# Neuromuscular connectomes across development reveal synaptic ordering rules

**DOI:** 10.1101/2021.09.20.460480

**Authors:** Yaron Meirovitch, Kai Kang, Ryan W. Draft, Elisa C. Pavarino, Maria Fernanda Henao Echeverri, Fuming Yang, Stephen G. Turney, Daniel R. Berger, Adi Peleg, Marta Montero-Crespo, Richard L. Schalek, Ju Lu, Jean Livet, Juan-Carlos Tapia, Jeff. W. Lichtman

## Abstract

In mammals, the connections between motor neurons and muscle fibers profoundly reorganize in the early postnatal period. To better understand this synaptic rewiring we traced out all the connectivity in muscles at successive ages in the mouse using serial section scanning electron microscopy in a muscle at birth and Brainbow-based and XFP-based fluorescent reconstructions in neonatal and older muscles respectively. Our data indicate that axons prune about 85% of their branches in the first two weeks of postnatal life, and that while much of this pruning leaves neuromuscular junctions with only one remaining axon (a ∼8-fold reduction), it also causes a ∼6-fold reduction in the number of muscle fibers that possess more than one neuromuscular junction. Unexpectedly, the simplification of the wiring diagram was not haphazard but rather was constrained by the tendency for neurons to maintain co-innervation the longest with other neurons based on their proximity in an abstract rank order. This synaptic ordering preference was even significant at birth when connectivity was the most overlapping but became more striking as development proceeded and was even obvious in the few adult muscle fibers that retained more than one axon at different neuromuscular junctions. Analysis of properties of muscle fibers sharing axons at developing ages and changes in the physical distance between neuromuscular junctions that were maintained in young versus older muscles suggests that the rank order of motor neurons is based on their relative similarity in activity patterns. This same ranking governs both the close-proximity synaptic competitions within neuromuscular junctions and the long-distance competitions that remove or maintain synapses millimeters apart meaning that all neuromuscular rewiring is based on the same global activity ordering rule. We think it is likely that this ranking is related to the ultimate recruitment order of motor axon activity as first described by (Henneman, 1957). Thus the emerging structure of neuromuscular circuitry is a product of its function: initial nearly all-to-all connectivity gives rise to a well-organized system of axons, allowing for the orderly recruitment of neurons during a smoothly graded behavior.

## INTRODUCTION

During a muscle action, motor neurons that generate little force, presumably because they innervate small numbers of muscle fibers (i.e., having small motor units), are activated first. If the force output is insufficient for the task at hand, progressively larger motor units are recruited. Such orderly recruitment of motor neurons, known as Henneman’s size principle, is a nearly universal property of motor action in vertebrates (Henneman, 1957; Henneman & Olson, 1965; Zajac & Faden, 1985). Although there are several theories, how this recruitment pattern comes into being remains unclear (Freund, 1983; Zucker, 1973). This problem is further complicated by the fact that motor units are larger in early postnatal development, and individual muscle fibers are multiply innervated due to the convergence of more than one motor axon at individual neuromuscular junctions (NMJs, also known as motor endplates). In adults, in contrast, each muscle fiber typically contains a single NMJ, generally located near the center of the fiber and contacted by only one motor axon. Thus, both the fan-out of individual motor axons and the fan-in of multiple axons decrease as animals are first using their muscles in postnatal life. In addition, the axon that remains at each NMJ establishes many new synaptic contacts there, more than compensating for the loss of other inputs (Sanes & Lichtman, 1999). In rodents this synapse elimination is typically completed in the first several weeks after birth. To the best of our knowledge, a link between the size principle and synapse elimination has not been made previously.

This change in connectivity over early development raises a number of questions including some about the size principle. 1) Are motor units distributed in size in early postnatal life or do motor units size differences emerge because of synaptic pruning? 2) Is developmental synapse elimination related in any way to the size principle? 3) What is the role, if any, of activity in establishing the size principle? The latter question arises because evidence suggests that the outcome of synapse elimination is based on activity differences between axons that innervate the same muscle fiber (Balice-Gordon & Lichtman, 1994; Buffelli et al., 2002; Busetto et al., 2000; Favero et al., 2012; W. A. Harris, 1981; Personius & Balice-Gordon, 2001; Wyatt & Balice-Gordon, 2003). Owing to the recruitment order, an individual motor neuron has an activity pattern that is most similar to the neuron recruited just before and just after it. We wondered if, as synapse elimination removes axons from NMJs, the axons with the most similar activity patterns would coexist at the same NMJ for longer than those with very different activity patterns.

All of these questions require knowing the complete set of neuromuscular connections for each and every motor neuron innervating a muscle. In a previous work, we reconstructed all the axons that innervated a small muscle in the adult mouse (Lu et al., 2009). In the present work, we performed the same type of dense reconstruction at multiple developmental ages. This connectomic approach was challenging because we needed to resort to serial electron microscopy for the youngest age studied (the day of birth, P0). At the end of the first postnatal week, we could use multicolor Brainbow mice (Livet et al., 2007; Tsuriel et al., 2015) with confocal microscopy. At later ages the preparation was large enough, so transgenic mice expressing a single type of fluorescent protein (XFP) (Feng et al., 2000) sufficed for the reconstruction as has been done previously (Lu et al., 2009).

Our results provide evidence that the size principle has its origins in the earliest stages of postnatal development and perhaps even earlier, but synapse elimination in postnatal life also plays a critical role. We found a hierarchy in the connectome among motor neurons. We believe that this hierarchy determines the order in which supernumerary axon terminals are removed from each NMJ. This organizational principle is compatible with the idea that synapse elimination is driven by differences in the activity patterns of motor neurons based on their place in the recruitment order. Lastly, we found evidence for a different kind of natural synapse elimination in postnatal life: the elimination of supernumerary endplates, which we ascribe to activity-based synapse elimination working over long distances.

## RESULTS

We compared the patterns of connectivity between lower motor neurons and muscle fibers in 31 individual muscles from 21 mice, ranging from newborn to adult. These datasets include serial-section electron microscopy (EM) analysis of the entire connectional map (connectome) of an interscutularis muscle at birth (postnatal day 0, P0; Fig. 1a), brainbow-based reconstructions of seven omohyoid and lumbrical muscle connectomes at P6-9 (Fig. 1b), and XFP-based reconstructions of two interscutularis muscle connectomes at P60 (Fig. 1c). For P3-5, P10-12, and P30, we reconstructed partial connectomes. As serial-section EM takes considerable time and effort, we only used it for the P0 reconstruction and obtained full connectomes at later ages with confocal microscopy instead. To aid in the segmentation of axons at P6-9, we bred brainbow mice and selected muscles with axons labeled with distinct colors (see Methods). In the adult, monochromatic fluorescent protein expression suffices for tracing all the axons as shown previously (Lu et al., 2009). We also analyzed the size of each muscle fiber in all of the data sets providing us with a way to link motor neuron properties to muscle fiber properties.

**Figure 1.**
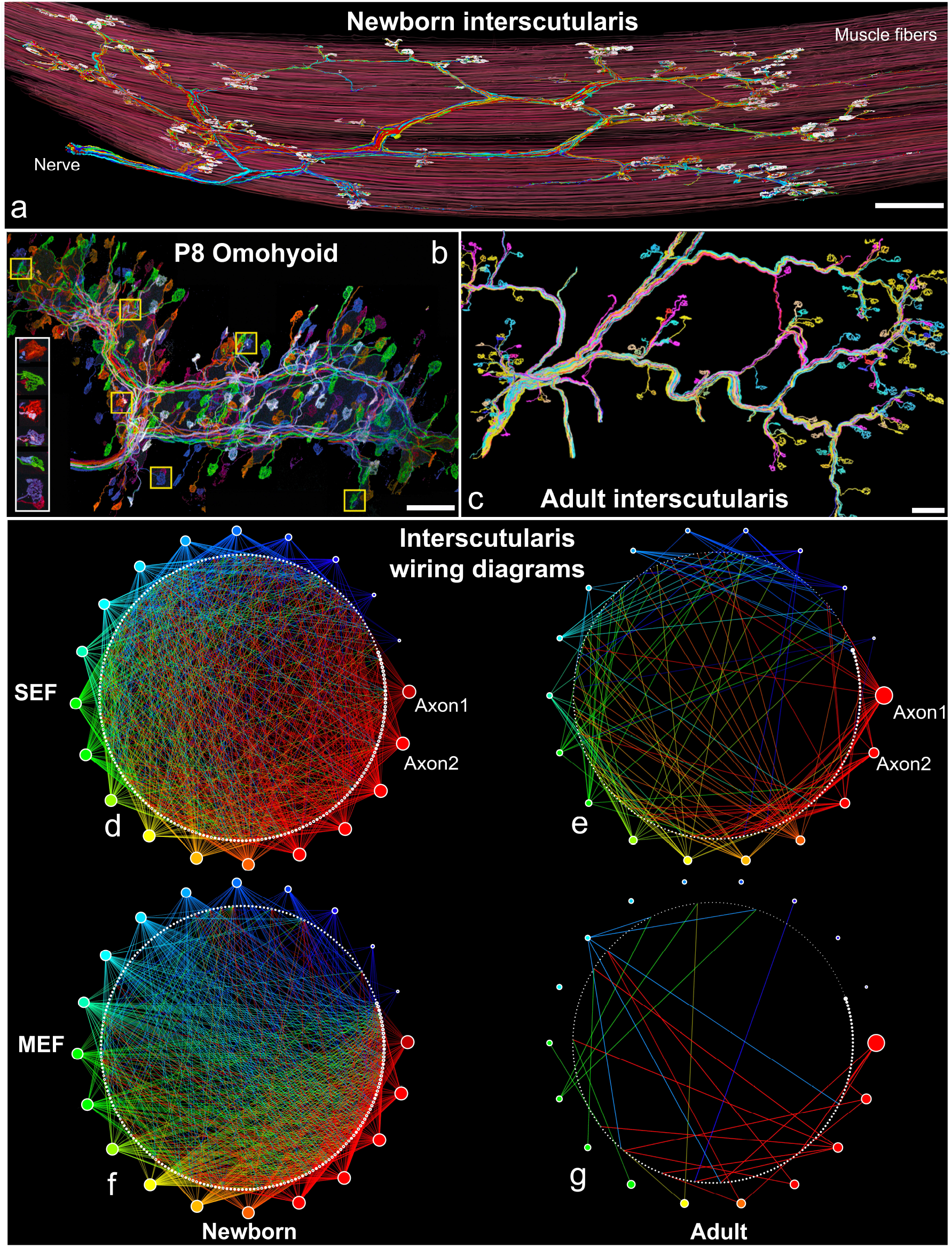
Muscle anatomy and wiring diagrams. **a**, A newborn interscutularis muscle and the innervating nerve reconstructed from serial-section electron microscopy images using machine learning-aided 3D tracing. Axons are colored according to their motor unit sizes (number of innervated muscle fibers). Red and blue for large and small motor units, respectively. Muscle fibers (red) and endplates (white) were individually reconstructed. **b**, The connectome of Brainbow-labeled axons innervating a P8 omohyoid muscle (Livet et al., 2007; Tsuriel et al., 2015). Boxed regions (white and yellow) depict typical multiply-innervated junctions. **c**, The connectome of XFP-labeled axons innervating an adult interscutularis muscle, reconstructed from high-resolution confocal microscopy images. **d-g**, Wiring diagrams show a comparison between newborn and adult connectomes demonstrating the massive axonal rewiring that takes place at these developmental ages. Colored circles represent motor axons (circle diameter proportional to the motor unit size). White circles represent muscle fibers (circle diameter proportional to fiber cross-sectional area). There are >1800 connections at birth but only ∼200 in adulthood. About 35% of the newborn muscle fibers possess multiple endplates, whereas ∼94% of the adult muscle fibers possess a single endplate. Connections made on single-endplate fibers (SEFs, d-e) and multiple-endplate fibers (MEFs, f-g) are shown separately. Scale bar: 100 μm.

### The newborn and adult interscutularis connectomes differ in several ways

The neuromuscular wiring diagram changes extensively in postnatal life (Bennett, 1983; Hall & Sanes, 1993; Purves & Lichtman, 1980; Sanes & Lichtman, 1999; Willshaw, 1981). To better understand the rules governing these changes, we compared the neuromuscular connectomes at birth and in adulthood (P30-P60; Fig. 1d-g). We found three major differences. First, nearly 100% of endplates in the newborn muscle received polyneuronal innervation, *i*.*e*., innervated by more than one motor axon. However, the number of converging inputs onto newborn endplates varied considerably (6.53 ± 2.17, range: 1-13; see also Fig. S3 and (Juan C. Tapia et al., 2012)). This ratio of convergence is greater than that reported by previous electrophysiological studies of newborn rodent muscles: for example, 2-4 terminals per endplate in E21-P0 rat diaphragm (A. J. Harris & McCaig, 1983; Redfern, 1970), 3.1-3.5 in P4 rat lumbrical muscle (Jones et al., 1987), and 2-6 in P0 rat extensor digitorum longus (EDL) (Balice-Gordon and Thompson, 1988). This discrepancy is consistent with the methodological limitations of measuring weak synapses by electrodes (Bennett & Pettigrew, 1974; Brown et al., 1976). By contrast, 100% of adult endplates were innervated by just a single axon. Second, 34.4% of the newborn muscle fibers possessed two or three endplates (multiple endplate fibers, MEFs), each of which received polyneuronal innervation. Twenty muscle fibers (27.6%) had two endplates and five fibers (6.7%) had three. The distance between endplates on the same muscle fiber was highly variable (510 ± 308 µm, range: 27.54 - 974.877 µm). By contrast, many fewer fibers in adult muscles had two endplates (7.7 ± 6.2, range: 1.4% - 18.1%, n = 5 muscles) (Fig. 1g, 4a). One adult P30 muscle had only 3 muscle fibers with multiple endplates and no adult muscle had any fiber possessing three endplates. Thus, during development both the number of axons per endplate (neuromuscular junction fan-in) and the number of endplates per muscle fiber decrease (Fig. 1d-g). Although it has been shown that acetylcholine receptor (AChR) plaques are removed from early prenatal muscle fibers (Sanes & Lichtman, 2001), our P0 data indicate that postnatally, wholesale endplate loss is also commonplace. Third, the motor unit size (i.e., the number of muscle fibers innervated by each motor axon) was far larger in the newborn: the average was 91.7 ± 32.4 fibers at P0 (range: 13 - 120, n = 20 motor units), but shrank to 14.1 ± 8 fibers at P60 (range: 3 - 39; n = 34 motor units from 2 connectomes) (Fig. 1d-g). In both the newborn and adult, motor unit sizes varied substantially. In adults, the size of a motor unit is thought to be related to the type of the constituent muscle fibers (e.g., fast vs. slow muscle fibers (Burke et al., 1971)). However, the variability in the size of the motor units observed in the newborn was unexpected, implying that even at birth motor axons are already “heterogeneous” (see Fig. S2). For example, the smallest motor units at birth clearly could not give rise to the largest motor units in the adults, because even before pruning they innervated fewer muscle fibers than the largest adult motor units (see Fig. 2b). Thus, a non-uniformity of motor units already exists prior to the phase of massive synapse elimination. This conclusion is buttressed by the finding that axons innervating the smallest motor units at P0 had the smallest calibers (measured at the nerve entrance, Fig. 2c), as they do at later ages.

**Figure 2.**
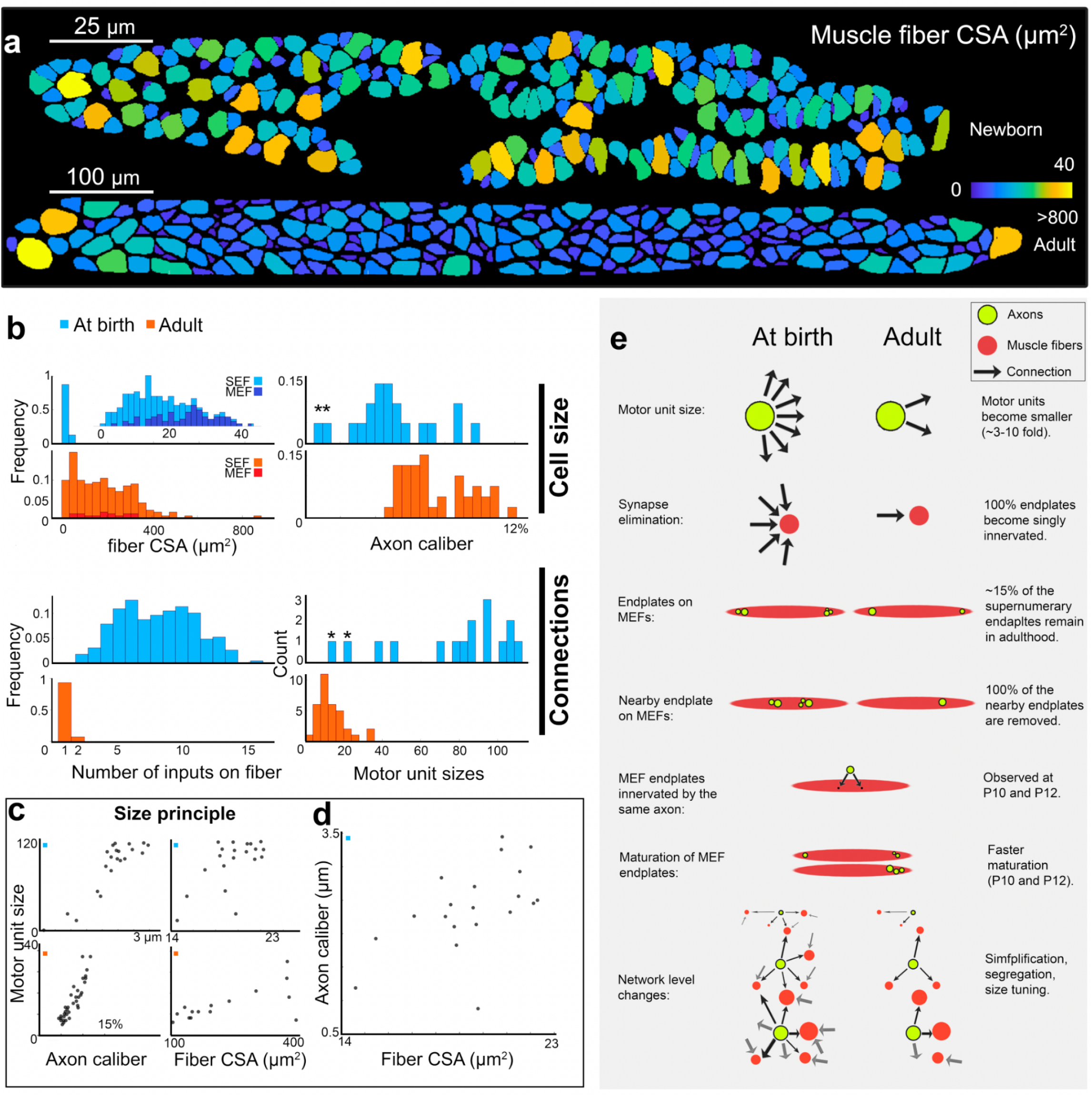
A size correlation between axons and their innervation targets from birth to adulthood. **a**, Transverse sections of a newborn and an adult interscutularis muscle. Individual muscle fibers colored by cross sectional area (CSA). The scale bar indicates the CSA distribution. **b**, Newborn muscle fibers are ∼10% the size of adult muscle fibers. At birth multiple-endplate fibers (MEF) are larger than single-endplate fibers (SEF); this size difference disappears in adulthood. The caliber of motor axons in the newborn shows a bigger distribution in sizes than that of adult axons. The number of distinct axonal inputs onto individual muscle fibers is highly variable (2-15) at birth and reduces to one or two inputs in the adult for SEFs and MEFs, respectively. The motor unit size drastically decreases from birth to adulthood and the shape of the distribution nearly flips. The two smallest neonatal motor units (marked by asterisks) are too small to mature into the large motor units in the adult. **c-d**. Axon calibers, motor unit sizes, and the average size of innervated muscle fibers are all correlated both at birth and in adulthood, suggesting that features of Henneman’s size principle (Henneman, 1957) start to emerge in perinatal life. **e**, Summary of the morphological changes observed in neuromuscular connectomes during postnatal development.

Despite the extensive synapse elimination (including the removal of entire endplates) from birth to adulthood, neither the total number of muscle fibers (P0: 222-270, adults: 205-254, n = 5 muscles each) nor the total number of motor axons (P0: 16-20, n = 3 nerves; adults: 16-18, n = 2 nerves) differed substantially across age. This similarity in the number of pre- and postsynaptic cells allowed us to directly compare the pattern of innervation from birth to adulthood. At birth, the larger motor units, the polyneuronal innervation of endplates, and the existence of multiple endplates on single muscle fibers together gave rise to a connectivity pattern that approached an all-to-all wiring diagram. Indeed, the total amount of actual connections between motor axons and muscle fibers (1833) was about 40% of the theoretical maximum (all-to-all with 240*20=4800 connections). In stark contrast, in adulthood only ∼6% of the potential connections were realized.

Although the neonatal neuromuscular connectivity exhibited extensive fan-in (convergence) and fan-out (divergence), motor axons did not randomly innervate muscle fibers. Rather, both motor axons and muscle fibers showed connectional selectivity even at birth. Five aspects of this non-randomness were evident. First, motor units were more variable in size than expected from a uniform random connectivity model (see Methods). In particular, the standard deviation of motor unit sizes, 32.4, is unlikely to arise from a random distribution of connectomes (*p*<10^−5^).

Such variability was manifested in the fact that 11 motor units each innervated at least 100 muscle fibers and 4 motor units each innervated only 13-53 muscle fibers, exhibiting a 9.2 fold size difference between the largest and the smallest motor units. Second, the number of axons converging onto single muscle fibers was highly variable (range: 3-16), inconsistent with a uniform random innervation model (*p*<10^−5^ using random connectomes with motor unit sizes the same as in the data). Third, the number of inputs per muscle fiber was significantly correlated with the cross-sectional area (CSA) of the muscle fiber (Fig. S1a; *p*<0.001). Fourth, the motor unit size and the average muscle fiber CSA among the muscle fibers innervated by a motor axon were correlated (*r*=0.629, *p*=0.0029). Larger motor units tended to innervate muscle fibers with larger CSAs (in total 10/20 motor units showed this preference; *p*<0.01 as in Fig. S1b; group selectivity compared against random connectomes was *p*<0.001; see Methods). Fifth, the motor axon caliber was correlated with both the motor unit size and the CSA of innervated muscle fibers (see Fig. 2c-d; Pearson correlation between axon caliber and motor unit size *r*=0.89, *p*<10^−7^; axon caliber and average muscle fiber CSA: *r*=0.56, *p*=0.011). In summary, axons with the smallest motor units were the thinnest, and tended to innervate the thinnest muscle fibers with the fewest converging inputs, and vice versa for the largest motor units.

This motor unit size-related innervation pattern is similar to that in adult muscles (see (Lu et al., 2009)), in which the motor unit size is correlated with the caliber of each motor axon at the entrance of the muscle. This correlation is part of the underlying reason for the orderly recruitment of progressively more powerful motor units (Henneman, 1957). The unexpected existence of size principle-like features at birth raises the possibility that axonal activity patterns are already “heterogeneous” in the newborn (see (Jones et al., 1987) for P3-P5). However, we could not detect other ultrastructural features that contrived with muscle fiber caliber such as sarcomere spacing, z-line thickness, and nuclear domain sizes, features that do differentiate muscle fiber types in adult muscles (Granzier et al., 1991; Tseng et al., 1994). Rather, these features were unimodally distributed among the muscle fibers. Nonetheless, newborn muscle is populated by a heterogeneous population of both motor axons and muscle fibers.

The heterogeneity in the neonatal innervation pattern prompted us to assess whether certain pairs of axons are more likely to share targets. We calculated the probabilities of observing the number of co-innervated muscle fibers between each pair of motor axons, under the null hypothesis that each axon innervates each muscle fiber with equal probability independent of other axons (Fig. 3/S5; see Methods). We found in the newborn connectome that many axonal pairs co-innervated significantly more muscle fibers than expected by the null model (2000 random connectomes; *p*<0.001, see Methods). Interestingly, some pairs exhibited significant avoidance of innervating the same muscle fibers (*p*<0.001). This organization of motor axons based on the targets they share, or avoid, indicates that the innervation pattern of axons is not random and rather reflects differences between motor neurons. However, learning what the overall pattern is, could not be discovered by this pairwise analysis and required looking at the entire network.

### Ordered pattern of innervation in newborn

In order to examine if the nonrandom innervation of muscle fibers by motor axons was part of a more global connectional scheme, we mapped the pairwise co-innervation pattern on a graph. In this graph each node represented one axon and each edge corresponded to a pair of axons co-innervating significantly more or fewer muscle fibers than expected by chance. We then weighted each edge by the degree to which the co-innervation deviated from the chance level. This approach allowed all axons to be normalized to the same scale independent of their motor unit sizes. To visualize this network, we embedded it into a plane using a force-directed graph drawing (Morgan et al., 2016). Axons that co-innervated significantly more muscle fibers than expected by chance were pulled together; conversely, axons that shared fewer muscle fibers than expected by chance were farther apart. This axonal network appeared elongated in one direction (Fig. 3a). To determine whether this global elongation could have emerged by chance, we simulated 500 random connectomes constrained by the same fan-in and fan-out statistics of motor axons and muscle fibers found at birth (i.e., the “configuration model”; (Newman, 2018)), and computed a metric of network elongation for each simulated connectome (see Methods). We discovered that the actual P0 connectome was more linear (elongated) than expected by chance (*p*<0.04). This elongation signifies that axonal preferences are ordered in a roughly linear way: axons that are adjacent in the order have preferences to co-innervate the same muscle fibers, axons that are distant in the order share little innervation to the same muscle fibers. This order appears to be a continuous set of preferences rather than two or more strictly selective cohorts - as would occur if there was a strict specificity between motor neurons of one type for muscle fibers of a corresponding type (Fig. S5). What neuronal property underlies this ordering will be discussed below.

**Figure 3.**
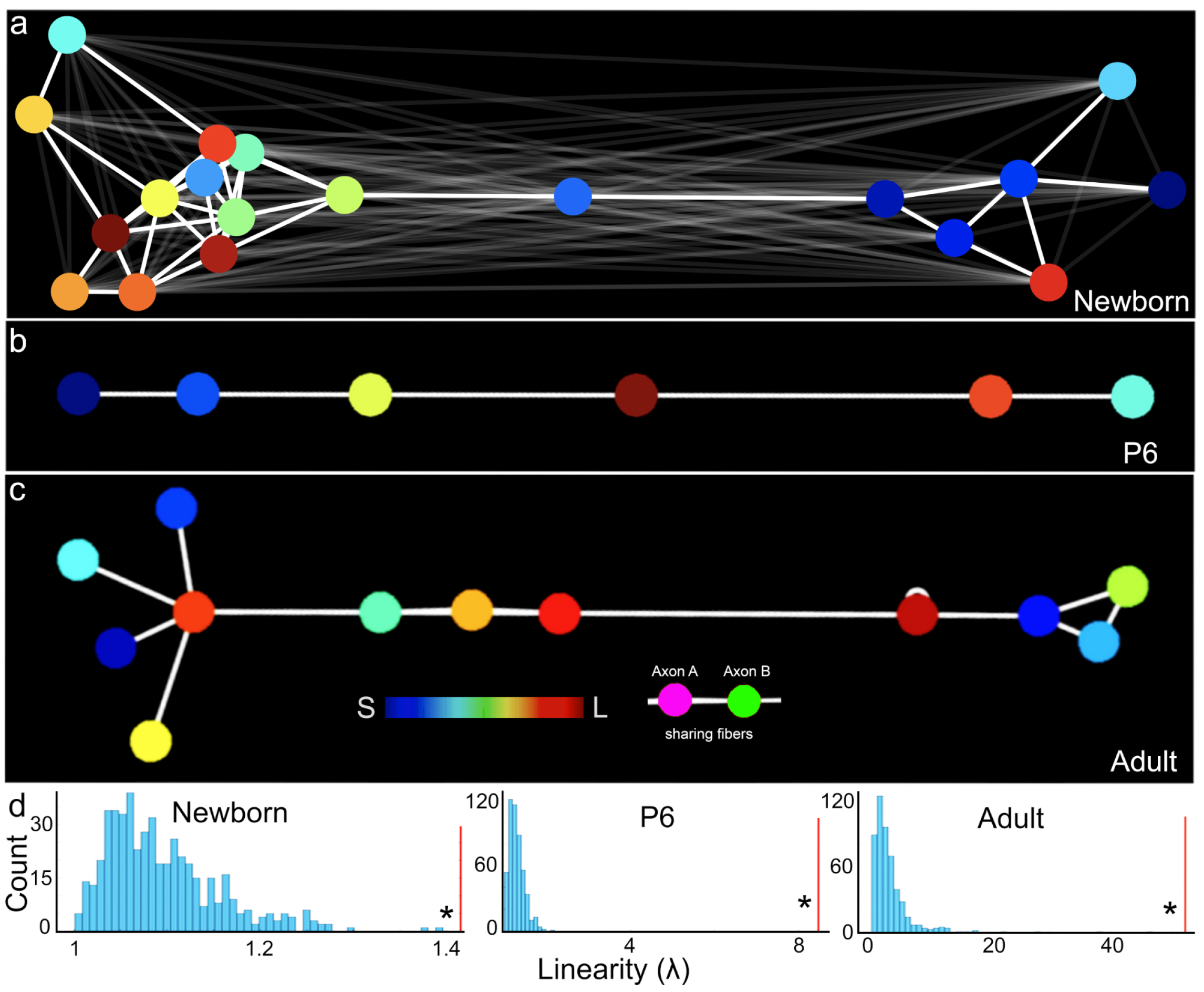
Linear order of axonal networks across maturation. **a-c**, The nodes in the graph representation correspond to axons whereas edges represent the number of shared targets between pairs of motor units. A pair of axons are connected by an edge if they co-innervate significantly more targets than expected by chance (a-b). The graph representation of adults obtained by analyzing MEFs connectivity (c). The edge length represents the likelihood to observe the actual number of shared targets under the assumption of independent, uniform random innervation (short edges for excessive shared innervation). Nodes are positioned by an optimized 2-dimensional embedding using a spring-force algorithm (Fruchterman & Reingold, 1991). The linearity within these axonal graphs (**λ**) was measured at P0 (n=1), P6-P9 (n=7), and P60 (n=2) by comparing its elongation (eccentricity of the scattered nodes) to that of random connectomes with the same number of axons, fibers and motor unit sizes. The ability to embed the networks in 2D and obtain a large elongation index indicates that axons are rankable such that an axon’s innervation proclivities were most similar to the axons on either side of it in the linear ranking. **d**, All three connectomes show elongation of the axonal network significantly beyond that of a random network (*p*<0.05). The red vertical lines indicate the elongation of the biological connectomes, compared against random size-matched connectomes (cyan).

To make this ordering more concrete we provide this example (Fig. S4a). Axon #8 shares a significant number of muscle fibers with axons #1 and #10, and both of them share significant numbers of muscle fibers with each other and with axon #2. In addition to these connections, axon #2 also shares a significant number of muscle fibers with 4 axons that are part of a central cluster of axons that mutually share many muscle fibers. On the north side of that cluster, axons #18 and #5 each shares a significant number of muscle fibers with several axons in the central cluster as well as with each other. However, these two axons (18 and 5) do not have significant shared connectivity with any axons on the south side of the central cluster. Both of them however, share a significant number of muscle fibers with axon #6, an axon that is not sharing a significant number of muscle fibers with any other axon, and hence is the northmost axon in this ordered array. This arrangement indicates that many motor axons have strong preferences to share muscle fibers with just a few other motor units, and those motor units in turn have strong preferences for a slightly shifted subset of motor units. We thus termed this elongation a “linear order of co-innervation”.

Furthermore, we found that axons that preferentially innervated muscle fibers with large CSA were located close to the middle of the linear order (i.e., where the central cluster of motor units resides; Fig. 3a and Fig. S4c-d). In contrast, axons with no preference for large muscle fibers were at the edges (poles) of the linear order (Fig. S4c-d). Most axons at the periphery sampled from both small and large muscle fibers, except one axon (#6) that appeared to be specific to thin muscle fibers. Thus motor axons that are specific to larger fibers share more targets with other axons of their kind.

One hypothesis that could provide an explanation for this ordering would be if motor units were topographically arranged so that the sharing was secondary to where in the muscle axons established synaptic branches. Although some small motor units were by virtue of their size unable to uniformly sample muscle fibers throughout the muscle, all of the larger motor units were widely distributed, and, as previously described in the adult, spatial segregation of motor units is not seen in this muscle in adults or in younger muscles ((Lu et al., 2009) and see below). Hence spatial maps do not explain this ordering.

### The adult neuromuscular connectivity cannot have developed from the newborn pattern by random synapse elimination

The innervation pattern of muscle fibers in the newborn indicates that the connectivity of motor axons is arranged in an ordered system to some degree. In addition, the largest motor units have the largest axon calibers and show a tendency to contact muscle fibers that also have the largest CSAs. These tendencies potentially are a prelude to the size principle observed in the adult muscle. Because the connectivity is much denser (∼8x) in newborns, we asked if the adult connectivity pattern could emerge from the neonatal connectome by random synaptic pruning or if it requires a more specific refinement. We found that the distributions of motor unit sizes in the newborn and in adults (n=2 and see also (Lu et al., 2009)) are not scaled versions of one another: the newborn has a right-skewed distribution of motor unit sizes (only few small units and several large motor units) whereas the adult has a distribution of motor units sizes that is skewed to the left (with only 1-2 large motor units) (see Fig. S6 for comparisons of the histograms). This result in and of itself does not resolve the role of neonatal pruning because some muscle fibers are already non-randomly innervated by more axons. We thus modeled the outcome of a gradual and random elimination of synapses from the newborn connectome to see whether it leads to the adult connectome. The simulated adult connectomes arising from random synaptic pruning have significantly different motor unit size ranges, mean sizes and variance (Fig. S7). Notably, the largest motor units in the adult connectomes (35 and 39 muscle fibers) are far larger than the random elimination model predicts (mean 20.09 ± 2.05 muscle fibers, estimated *p*<0.01; 0/1000 occurrences of null hypothesis in randomly produced developing connectomes with the starting point taken from the P0 connectome). Hence, although the wiring between motor axons and muscle fibers shows some level of specificity at birth, this pattern is not sufficient to generate the adult connectome.

### The linear order after birth

The above makes the case that pruning cannot be random but does not clarify what rules govern which connections are removed and which are maintained. One possibility is that co-innervation persists longer among motor axons that have a similar rank order. To examine this idea, we reconstructed neuromuscular connectomes at P6-P9, when many connections have been pruned but some muscle fibers remain multiply innervated. At this slightly older age, muscles are much larger, rendering serial EM reconstruction impractical. We therefore opted for the brainbow transgenic labeling strategy with confocal imaging, which in the larger and less complicated older muscles provides sufficient resolution to map all connections. The expression of the brainbow transgenes under the Thy-1 promoter was weak in rostral muscles, including the interscutularis at P6, so we examined instead several small and more caudally-located muscles in the neck (omohyoid) and the forepaw (lumbrical) where expression was adequate. We reconstructed the connectomes of four omohyoid (OM 1-4) and three lumbrical muscles (Lum 1-3), and generated the spring models of these networks as above. An obvious linear order was evident in six of the seven muscles (all omohyoid and 2 of 3 lumbrical muscles). Moreover, one muscle presented a perfect linear order (Fig. 3b and Fig. 5 and Discussion), suggesting that it is not a unique property of the interscutularis muscle and that the ordering becomes more robust after some pruning has taken place. The one muscle that did not show a linear order (Lum 2) was at an advanced stage of synapse elimination, with 73.48% of the endplates already singly innervated so there were gaps in the ordering. However, in that muscle there was still strong evidence that axons that continued to share muscle fibers did so in a way that was unlikely to have occurred by chance (p<0.01; Monte Carlo test measuring the total amount of sharing). Hence, we conclude that the linear order does not weaken after the first stage of the massive postnatal synapse elimination. This indicates that motor axon synapses are progressively eliminated according to innervation tendencies that already appear at birth. In addition, these findings suggest that motor axon synapses are eliminated based on developmental properties of motor neurons and muscle fibers, rather than based on purely stochastic processes. Moreover this order is unrelated to the topography of axon branches, ruling out a mechanical explanation for this specificity. This also suggests that the set of muscle fibers co-innervated by two axons will maintain this co-innervation for a long period while other axons innervating the same muscle fibers will retract earlier. This conjecture raises the possibility that the linear order might still be observed into adulthood, as manifested by the identity of the axons that share multiple endplates on the same muscle fiber.

### Evidence of ordered innervation in adult muscles: multiple endplates on muscle fibers

At developmental ages, we inferred graded specificities of motor units for muscle fibers by virtue of the extensive co-innervation of muscle fibers by shifting subsets of motor neurons. As most adult muscle fibers are innervated by single axons at their solitary neuromuscular junctions, one cannot directly relate the developmental target-sharing to their adult state of innervation. However, as mentioned above the adult interscutularis muscle retains a small number of MEFs (∼15% of the number of MEFs at birth; Fig. 4a). Hence, developmental synapse elimination mechanisms both reduce the number of motor axons per endplate and the number of endplates per muscle fiber. We examined whether the remaining MEFs provide a link to the developmental ordering phenomenon and the ultimate adult pattern of innervation (i.e., the size principle). We used confocal microscopy to image two interscutularis muscles from adult YFP-16 mice, which express yellow fluorescent protein in all motor axons (see Methods), and reconstructed their full connectomes including the axons, the muscle fibers, and NMJs.

**Figure 4.**
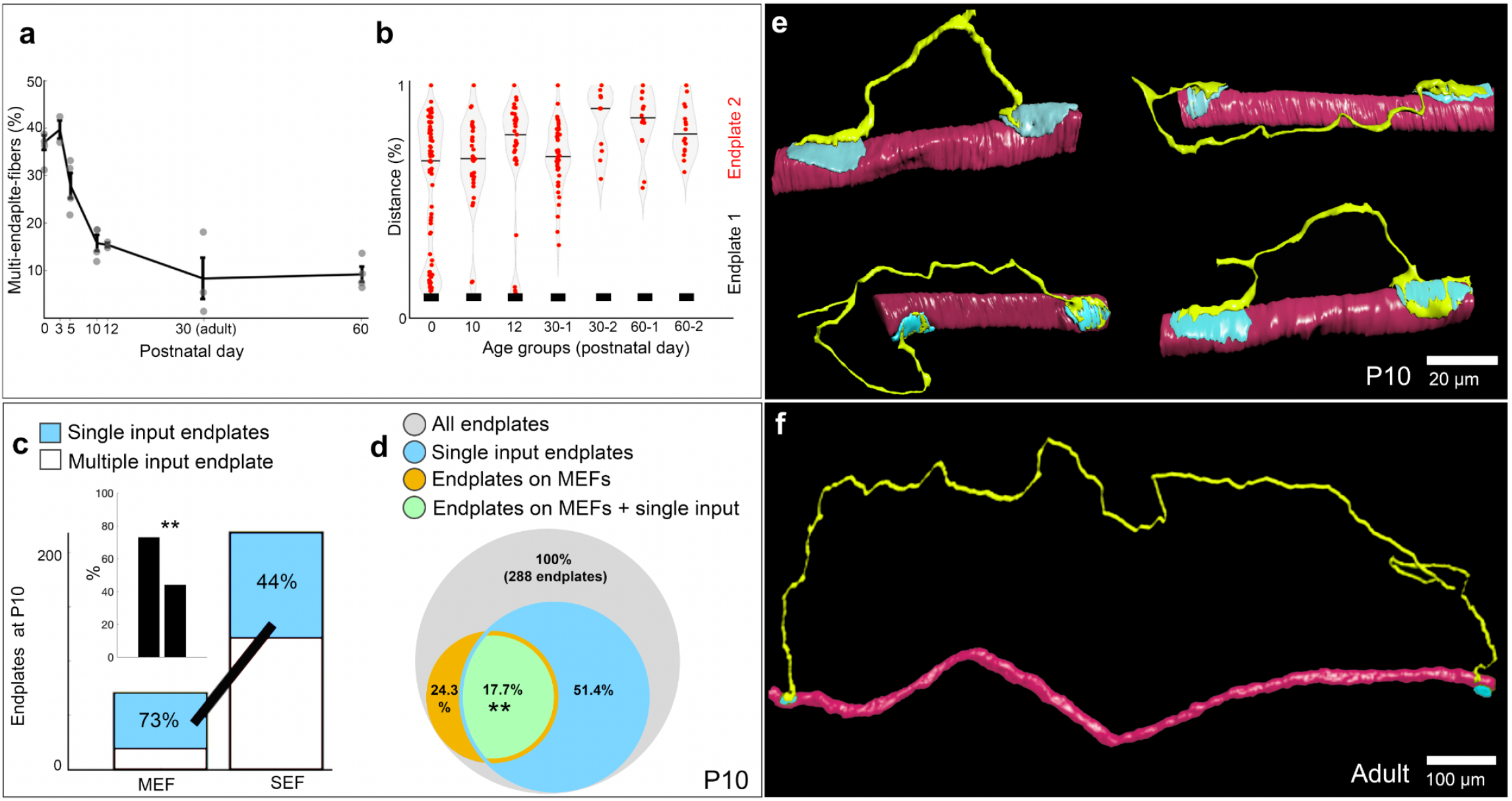
Action at a distance: distance dependence and axon identity dependency on the fate of elimination between paired endplates. **a**, Removal of supernumerary endplates across development. **b**, Distance dependence in the elimination of supernumerary endplates across maturation. **c-d**, In an intermediate developmental stage (P10), the endplates on MEFs are closer to becoming singly innervated than endplates on SEFs, which are more likely to continue to be multiply innervated (c) while the conjunction of an endplate to be both on MEF and singly innervated occurs significantly more often that expected by chance (d). This suggests that synapse elimination in endplates on MEFs is more advanced than that on SEFs. **e**, The few pairs of endplates that are nearby at P10 are innervated by the same axon. The distance dependence (a-b), the faster elimination on MEFs (c-d), the mono-neuronal innervation of nearby endplates at P10 (e), the presence of an order between the axons innervating pairs of junctions on MEFs, all suggest that the presence of an additional endplate on a fiber, and the specific axons innervating the two junctions, affect the fate of additional endplates on the muscle fiber. We conjecture that collision of action potentials may be the mechanism that allows remote endplates to develop independently of each other and coexist in adulthood (f).

The two muscles were similar: muscle A had 235 fibers and was innervated by 18 axons; muscle B had 217 fibers and was innervated by 16 axons. There were 13 MEFs in muscle A (5.53%) and 16 in muscle B (7.37%). In total, 26 endplates in muscle A (10.48%) and 32 endplates in muscle B (13.73%) belonged to MEFs. These endplates were singly innervated as were the solitary endplates on SEFs. In muscle A, 12/18 axons (66.67%) innervated at least one endplate on MEFs; in muscle B the fraction was 43% (7/16 axons). The connectivity to the MEF muscle fibers was unlikely to have occurred by chance. For example, on 4 fibers, the same axon innervated both NMJs on the fiber. Furthermore, there were 6 instances in which the same pair of axons co-innervated more than one MEF. In muscle A, only two MEFs were mono-neuronally innervated, which is not an unlikely event to occur by chance, but as we show below these pairs of co-innervating axons were ordered linearly, consistently with connectomes at younger ages. In muscle B, 6 MEFs received identical axonal inputs (p<0.032). There were even two axons each of which innervated two MEFs at both endplates. This strong tendency for axon pairs to co-innervate endplates on adult MEFs is consistent with the tendency of axons to repeatedly co-innervate the same endplates at younger ages (as described above). All of these suggest that the identities of the axons in the adult that are co-innervating the same muscle fibers are associated with each other and are not drawn from a random distribution of axon pairs. As described earlier, all of these patterns are central features of the connectivity at birth and in P6-P9.

The extra sharing as indicated on adult MEFs was related to an ordered system of innervation that was also reminiscent of the patterns of innervation seen at younger ages. In muscle A (Fig. 3c), the MEFs were connected by axons in a linear order that is unlikely to occur by chance (p<0.001). As in the newborn linear order, we found that also in the adult, the order of the neurons in the linear order was compatible with the tendency of some axons to specifically innervate, more often, fibers with larger CSA. First, the 5 motor units encompassing the thinnest muscle fibers (which was a subset of the 8 smallest motor units) did not innervate any MEF (out of the total of 6 motor units that did not innervate MEFs). Second, among the six axons that preferentially innervated small fibers (rank sum test, p<0.05), five did not innervate any MEF and the sixth was located at the periphery of the linear order. The two axons that preferentially innervated the large muscle fibers were located in the center of the linear order, and were the two largest motor units (adjacent to each other in Fig. 3c). In muscle B (Fig. S8) the connectivity pattern is also highly unlikely to have occurred by chance. As already mentioned, the same pairs of motor units share more than one MEF. Moreover, two axons concentrated their innervation on MEFs to a greater degree than expected by chance: Axon 1 innervated 9 out of 13 MEFs (69.23%) and a total of 10 endplates on MEFs (38.46% of the 26 MEF endplates) (p<0.05). Axon 2 innervated 5 MEFs (38.46%) and a total of 7 endplates on MEFs (26.92% of the MEF endplates) (p<0.03). This repeated innervation of endplates by 7 axons (p<0.005; see MC in Methods) indicates, consistent with the linear order in the other adult muscle (A) and the order in younger ages, that the innervation pattern is structured from birth (dense connectivity) throughout maturation during the phase of massive synapse elimination and in adulthood (sparse connectivity) when some fibers still maintain multiple synapses.

This non-random connectivity implies that the MEF-innervating axons have something in common. However, we found no evidence that MEFs constitute a distinct group. For example, the caliber of an adult muscle fiber is a strong indicator of its functional type (Frontera & Ochala, 2015). However, adult MEFs exhibited a broad and uniform distribution of CSAs comparable to that of SEFs (bootstrapping test, p>0.1 in Muscle A). At birth, MEF fibers were thicker (p<0.01), but their sarcomere structures such as z-line thickness and sarcomere spacing were not distinct from that of SEFs (Fig. S11). The myonuclear domains on MEFs and SEFs were not significantly different at birth either. Moreover, the motor unit sizes of axons innervating MEFs (12/18 axons in muscle A and 7/16 axons in muscle B) were what we would expect to occur by chance (mean in Muscle A 16.1 endplates, p>0.1; mean in Muscle B 18.3 endplates, p>0.1, accounting for the centrality of the muscle; see Methods). Finally, the proportion of MEFs in five adult interscutaris muscles varied considerably: one muscle had only 3 MEFs (1.4% of the muscle fibers) and another had 46 (18.1% of the muscle fibers). Such a large variability suggests that, although interscutularis MEFs are innervated non-randomly, they are not essential for proper muscle function, and therefore unlikely to belong to one type (see Discussion).

### NMJ elimination from MEFs

The non-random connectivity in adult MEFs shares features with the innervation pattern at earlier stages, providing a potential link between the developmental ordering phenomenon and the adult pattern of innervation (i.e., the size principle). We thus wondered if the simplicity of MEFs could give us insights into the mechanism that foments, or in some cases prevents, synapse elimination. As the newborn muscle contains ∼6-fold more MEFs than adult muscles (see above), endplate loss is evident during the period of synapse elimination. We found that endplate loss strongly depends on the inter-junctional distance. In the newborn muscle there were 29 pairs of endplates (∼37.6%) that were separated by less than 50% of the maximal inter-junctional distance (the maximal distance between the pair of endplates on a MEF, 0.975 mm). However, in adult muscles we never observed a pair of endplates on the same fiber that were separated by less than 50% of maximal inter-junctional distance (∼1.88 mm). Rather, all the adult multiple-endplate pairs on muscles A and B were at least 0.965 mm apart (Muscle B). This distance dependence was highly significant (means differed across developmental stages; p<0.0001) and the distance between the closest endplates increased significantly across maturation: between newborn (P0) and all other developmental ages P10, P12, P30, and P60 (Fig. 4b; rank sum of the mean of the left side of the distribution; largest p-value p<0.025 between P0 and P10), and between newborn and intermediate stage (P10 and P12) (p<0.001), intermediate stage and adults (p<0.02) and newborn and adults (p<10^−9^). The loss of the nearby endplate pairs occurred gradually during the first weeks of postnatal life. Importantly, however, distance did not guarantee that both endplates would be maintained on the adult muscle fiber. Although more than 20% of newborn muscle fibers possessed endplates that were at least 0.5 mm apart (more than 50% of the maximal inter-junctional distance), MEFs accounted for a small fraction (∼6%) of adult muscle fibers, implying that most of the distant endplate pairs at birth had lost one endplate by adulthood. Interestingly, we observed at an intermediate age (P10) 4 pairs of nearby endplates on MEFs. In each case, the same axon innervated both endplates (Fig. 4e); nevertheless, we anticipate that one of the two endplates would be eliminated eventually because no close endplates were seen in any adult MEF. We also found 4 adult MEFs in which the same axon innervated both endplates (albeit they were much farther apart and presumably would be stably maintained throughout life). We therefore infer that endplate elimination requires action at a distance: in some manner, the presence of one endplate decreases the sustainability of another endplate on the same fiber, especially, but not exclusively, if the latter is nearby. However, we found no evidence that a muscle fiber with a single endplate ever loses it, which argues that endplate elimination, just as intra-junctional synapse elimination, is a competitive process: the fate of an endplate is related to the presence and position of another endplate on the same muscle fiber. This is analogous to the situation that the fate of one axon at a multiply-innervated endplate is related to the presence and position of other axons at the same junction (Turney & Lichtman, 2012). We also noted that endplates on MEFs became singly innervated earlier in development than those on SEFs. At P10, 73% of the endplates on MEFs were singly innervated, whereas a significantly smaller fraction (44%) on SEFs were singly innervated (p<10^−8^; Fig. 4c). As synapse elimination is widely believed to be activity-dependent (Balice-Gordon & Lichtman, 1994; W. A. Harris, 1981; Meister et al., 1991; Personius & Balice-Gordon, 2001; Stryker & Harris, 1986; Wyatt & Balice-Gordon, 2003), when two endplates coexist on the same muscle fiber, the presence of inter-junctional sources of activity in addition to intra-junctional competition may accelerate synapse elimination.

One possible mediator of the action at a distance that leads to endplate elimination is the activity in the postsynaptic muscle fiber. Previous studies suggest that synaptic sites not activated by local neurotransmitter release are destabilized by depolarization from another source (Balice-Gordon & Lichtman, 1994; Lichtman et al., 1985). However, we did observe a few MEFs in the adult interscutularis muscle, which suggests that under certain circumstances action potentials (APs) elicited at one endplate do not destabilize the other endplate. Unlike intra-junctional synapse elimination which removes all but one axon, the complete loss of an endplate removes all the axons from that site. This only occurs if an additional endplate on the fiber is maintained. In this sense the interjunctional endplate loss is competitive because the survival of one endplate is adversely affected by the presence of another on the same muscle fiber. As stated above, propagating APs could be the destabilizing factor. However, the destabilizing effect must be blocked in certain circumstances as some muscle fibers stably maintain two endplates in adulthood.

One potential mechanism of blocking competition is the collision and reciprocal annihilation of two APs propagating towards each other in the muscle fiber (Bernhard Katz & Kuffler, 1941; Lateva et al., 2002, 2010; McComas et al., 1984). AP annihilation occurs because following the propagating depolarization is the absolute refractory repolarizing wave, in which nearly all voltage-gated sodium channels are inactivated. This refractory condition prevents one AP from passing through another along an axon or, more relevant here, a muscle fiber. To better understand the coexistence of a subset of endplate pairs on MEFs, we considered the condition for AP collision to occur in a muscle fiber (see Methods, Fig. S9, and Table 1). If an AP arrives at the presynaptic terminal, it elicits an AP in the muscle fiber with a random synaptic delay. Muscular AP collision occurs when AP onsets differ by less than the time it takes for an AP to propagate from one NMJ to the other. Using values of axonal and muscular conduction velocity taken from previous studies, we applied the model to the MEFs we observed in several ages. First, in P10 muscles we found MEFs with one axon innervating both endplates that were close to each other (41-72 µm), meaning that a muscular AP only takes 28-48 μs to propagate from one endplate to the other. The path length difference between axonal branches was likewise small (13-73 µm), which translates into a difference in presynaptic AP arrival time of 21-118 μs. With the standard deviation of the synaptic delay being ∼ 0.25 ms, we estimated that the probability of AP collision was 10% or less (see Table 1 for details). In addition, axonal branches of small caliber may exhibit presynaptic blockage of APs (Krnjević & Miledi, 1959), which further reduces the probability of collision. Hence, even though the two endplates were nearby and innervated by two axonal branches similar in length and caliber, one endplate could still send an AP to destabilize the other, as the temporal relationship between APs is dominated by the synaptic delay in this case.

**Table 1.** Prediction of collision of postsynaptic AP in developmental multiple-endplate fibers with pairs of nearby endplates. The distance between an axon terminal and its nearest branching point (L1 and L2) and the distance between endplates (L) along the muscle fiber (MF) were estimated by reconstructing and skeletonizing axons and muscle fibers in VAST (Berger et al., 2018). The axon conduction velocities (CV-Axons) near terminals were obtained from (B. Katz & Miledi, 1965a), and see also (B. Katz & Miledi, 1965b), using properties of axons of similar diameter. Due to possible differences in the cable properties between the studies, the maximal experimentally observed axon terminal CV was used (in favor of collision). We assume that the standard deviation in the synaptic delay time is 0.25 ms, in line with (Katz & Miledi 1964). The model predicts that postsynaptic AP collision may occur only with very low probability at P10, which may suffice to induce interjunctional elimination.

| P10 MEF | CV-MFs | CV-Axon | Axonal branch length diff (L2-L1) | MF length (L) | AP propagation time (NMJ1 to NMJ2) | Activity onset NMJ2 | Collision prob. | margin - time diff |
| --- | --- | --- | --- | --- | --- | --- | --- | --- |
| | (m/s) | (m/s) | (μm) | (μm) | (ms) | (ms) | | (ms) |
| 1 | 1.5 | 0.62 | 19 | 41.3 | 0.028 | 0.031 | 0.062 | 0.003 |
| 2 | 1.5 | 0.62 | 40 | 59.3 | 0.040 | 0.065 | 0.088 | 0.025 |
| 3 | 1.5 | 0.62 | 73 | 72.3 | 0.048 | 0.118 | 0.103 | 0.070 |
| 4 | 1.5 | 0.62 | 13 | 65.6 | 0.044 | 0.021 | 0.098 | -0.023 |

**Table 2.** Muscle samples studied. Inventory of all muscle samples examined. The age (postnatal days after birth), total number of muscle fibers, NMJs, and percentage of MEFs per sample are listed. EM= sample used for electron microscopy; YFP= XFP sample used for confocal microscopy.

|  | Day | Fibers | NMJs | % |
| --- | --- | --- | --- | --- |
| P0-1 (EM) | 0.5 | 188 | 262 | 39.4 |
| P1yfp1 | 1 | 245 | 341 | 39.2 |
| P1yfp2 | 1 | 222 | 309 | 39.2 |
| P1yfp4 | 1 | 234 | 307 | 31.2 |
| P1yfp2 | 1 | 248 | 344 | 38.7 |
| P1yfp5 | 1 | 270 | 368 | 36.3 |
| P3-1 | 3 | 250 | 356 | 42.4 |
| P3-2 | 3 | 230 | 315 | 37.0 |
| P5-1 | 5 | 262 | 328 | 25.2 |
| P5-2 | 5 | 254 | 334 | 31.5 |
| P5-3 | 5 | 268 | 357 | 33.2 |
| P5-4 | 5 | 254 | 309 | 21.7 |
| P10-1 | 10 | 312 | 370 | 18.6 |
| P10-2 | 10 | 252 | 287 | 13.9 |
| P10-3 | 10 | 253 | 300 | 18.6 |
| P10-4 | 10 | 252 | 282 | 11.9 |
| P12-1 | 12 | 264 | 303 | 14.8 |
| P12-2 | 12 | 244 | 283 | 16.0 |
| P30-1 | 30 | 254 | 300 | 18.1 |
| P30-2 | 30 | 205 | 216 | 5.4 |
| P30-3 | 30 | 220 | 223 | 1.4 |
| YFP1 | 60 | 219 | 233 | 6.4 |
| YFP2 | 60 | 231 | 248 | 7.4 |

Hence, endplates that are separated by less than 100 µm will not be stable even if the innervating axons originate from the same neuron or are synchronous (Fig. S9). By contrast, some MEFs persist into adulthood, with endplates separated by at least 1 mm. For example, the axon in Figure 4f has two branches with a path length difference of 1.861 mm (2.515 mm - 0.654 mm), corresponding to a 0.1241 ms difference in AP arrival time at the NMJs (assuming 15 m/s conduction velocity in myelinated axons). The postsynaptic sites are 2.211 mm apart along the muscle fiber, corresponding to 1.474 ms of propagation time. In this scenario, the large propagation time relative to the presynaptic arrival time difference, and to the standard deviation of the synaptic delay, almost guarantees AP collision (Fig. S9). This dichotomy is a possible explanation for the emergence of the axonal order in the network of axons co-innervating fibers with multiple endplates (Fig. 3c) and implies that endplate elimination is also based on the relative activity patterns of the axons at each site.

## DISCUSSION

This study aims to better understand the extensive re-wiring of the peripheral neuromuscular circuit during early postnatal life. We reconstructed full neuromuscular connectomes at birth, at one week of age, and in adults, as well as partial connectomes at other developmental stages. The hope was that this description would offer clues into the mysterious underlying processes that lead to an orderly recruitment of motor units, known as Henneman’s size principle (Henneman, 1957), which, from small to large, allows each recruited axon to add a discrete tension step owing to the complete lack of co-innervation at the same NMJ by multiple axons. This non-overlapping partitioning of muscle fibers into motor units is an emergent property of development, because at birth motor units are much larger, and each NMJ may be co-innervated by 10 or more axons (Callaway et al., 1987; Juan C. Tapia et al., 2012). We believe that comparing wiring diagrams across ages, while painstaking, is informative. Indeed a recent longitudinal connectomic study on invertebrate development shows that the fine structure of a wiring diagram can be understood through the lens of nervous system maturation (Witvliet et al., 2021).

This connectomic analysis of the neuromuscular system led to a number of novel findings. First, the vast majority of neonatal motor units were substantially larger than any adult motor unit, and none of the axons was the sole input to any muscle fiber at birth. This implies that all classes of motor neurons projecting to a muscle undergo substantial axonal pruning. Second, despite the fact that motor units at birth were several fold larger than in adults, both were broadly distributed in size: even at birth there were a number of relatively small motor units, a few intermediate-size motor units, and several gigantic ones innervating about half of the muscle fibers. Third, there was a correlation between the size of the muscle fibers (cross sectional area) and the size of the motor units that provided innervation to them. This correlation between motor unit size and muscle fiber diameter was similar to the relationship between large motor units and the largest fast twitch muscle fibers in adults (Hämäläinen & Pette, 1993; Henneman & Olson, 1965; Zajac & Faden, 1985). The identity of motor units (by size) allowed us also to realize that the cohort of axons that shared innervation at the same neuromuscular junctions was not a random subset of all the axons, but that axons of similar motor unit size tended to share muscle fibers more often than expected in a random model. All of this could suggest that muscle fiber types were already in existence at birth and classes of axons strictly confined innervation to one of the classes, as is the case in adults. However, the tendency of axons to co-innervate particular muscle fibers, or not, appeared to be graded rather than binary. Large motor units did innervate large muscle fibers but this was a tendency not a strict rule. Moreover, the innervation preferences of axons could be ranked in a linear order such that an axon’s innervation proclivities were most similar to the axons on either side of it in the linear ranking. Small motor units were close to other small motor units in the order and large motor units were near other large motor units. But because there was considerable overlap of different kinds of motor units at each neuromuscular junction the ranking was subtle to ferret out at the earliest stage we looked at (see however below). This overlap among motor units could mean that the size principle emerges by gradual pruning of axon branches to parcellate muscle fibers into smaller and smaller groups until each group is part of only one axon’s motor unit (see more below). Finally, we found that many muscle fibers (>30%) possessed two or rarely three neuromuscular junction sites (endplates). Other studies have made note of multiple-endplate muscle fibers, especially in facial and neck muscles (Agduhr, 1919; Bendiksen et al., 1981; Brown et al., 1976; Cattell, 1928; Duxson & Sheard, 1995; Happak et al., 1997; Hunt & Kuffler, 1954; Iwasaki, 1957; Jarcho et al., 1952; Bernhard Katz & Kuffler, 1941; Lateva et al., 2002; Périé et al., 1997; Rossi & Cortesina, 1965; Sandmann, 1885; Walker, 1961; Zenker et al., 1990). The interscutularis muscle is also a neck muscle. Interestingly the axons that innervated the pair of neuromuscular junctions on a fiber were not always identical, further suggesting that strict chemospecificity between motor axons and muscle fibers was not likely the cause of the tendencies of multiple motor units to share more muscle fibers than expected by chance which we discuss below.

The ordering of motor units was subtle at birth because many axons co-innervated each NMJ. At P6-9, however, the remaining multiply-innervated endplates mostly received inputs from only two axons, so this ordering became more obvious. Using brainbow technology (Livet et al., 2007; Tsuriel et al., 2015) to obtain connectomes in P6-9 muscles, we found a striking pattern of innervation, in which axons showed a gradient of preferences that in many muscles could be arranged into a nearly perfect linear order, such that the axon at one extreme shared muscle fibers mostly with the axon next in “rank” and the same axon co-innervated neuromuscular junctions progressively more rarely with axons farther away in the axon ranking (Fig. 5). Axonal interactions at sites of neuromuscular junctions were therefore akin to the interaction of neighbors along a street: most interactions occur with immediately adjacent neighbors, less so with neighbors a house away, and with progressively fewer interactions with neighbors further down the block. However, unlike a street, this linear order was abstract: it was not explained by topographic features of where the synapses were located. To the best of our ability we were unable to map the linear order to a motor map in the muscle; nearly all axons distributed their branches over much of the muscle and the center of gravity of each motor unit was not shifted in a way that could explain the ordering.

**Figure 5.**
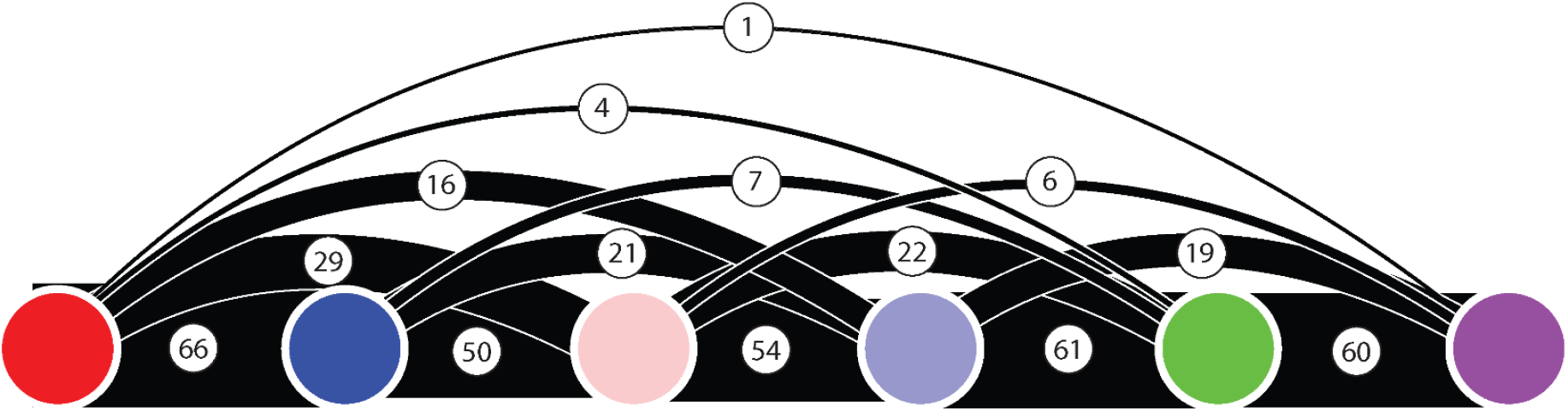
The linear order of connectivity at a P6 neck muscle based on brainbow labeling. This small muscle has only 6 motor units (represented by the colored circles). The number of muscle fibers co-innervated by a pair of motor neurons is given above the line connecting them (and the edge thickness). For example, the leftmost red motor unit co-innervates 66 muscle fibers with the blue motor unit, 29 muscle fibers with the pink motor unit, 16 muscle fibers with the lavender motor unit, 4 muscle fibers with the green motor unit and only one muscle fiber with the purple motor unit. Such co-innervation gradient is recapitulated for each axon. Remarkably, the same order (red, blue, pink, lavender, green, purple) describes the gradient of connectivities throughout the entire connectome. This order is not represented by a topographic map in the muscle.

Because of the topographic variability of motor units within the same muscle in different animals ((Lu et al., 2009) and confirmed here), one attractive idea is that the linear order is an emergent property explained by the similarities or differences in neural activity patterns among the cohort of axons innervating a muscle in a more or less random fashion. Remarkably, a similar ordering of preferences motif was described by Purves and colleagues as the laws of “contiguity” and “segmental dominance” in the spinal cord innervation of ganglion cells in the rodent sympathetic chain (Lichtman et al., 1979, 1980). In that system, preganglionic innervation from different but overlapping sets of spinal cord segments are linearly ordered in their innervation of cohorts of ganglion cells that project to different regions of the body. In both systems no topography was evident in the location of postsynaptic cells that shared the same subsets of axons (Lichtman et al., 1979, 1980). Similarly, we found no evidence of topography in the location of shared connections in the developing connectome (Fig. S15). Because reinnervation recapitulates the pattern of innervation the authors thought chemospecificity was the likely explanation for this pattern (Njå & Purves, 1977). In muscle we do not know if the origin of the ranking is also the position of neurons along the rostral-caudal axis of the spinal cord. Whatever its cause, its presence raises the possibility that this ordered overlapping connectivity is related to the emergence of the ordered recruitment of motor neurons in the size principle. We had assumed that making this link would be challenging, given that adult neuromuscular junctions are singly innervated, not allowing us to know which of the adult axons were sharing muscle fibers at earlier ages. But surprisingly a remnant of multiple innervation remained on a small number of adult muscle fibers in the form of the inputs to the few muscle fibers that maintained two neuromuscular junctions. These co-innervated adult muscle fibers showed preferences that were compatible with a linear order of specificity and thus provided a connection between the adult pattern of innervation and the linear order present at developmental ages.

One fundamental question is whether these preferences in muscle are based on chemical matching between motor axons and muscle fibers, or rather due to similarities in the timing or amount of activity. There are arguments supporting both ideas. As already mentioned, in the adult rodent superior cervical ganglion there is a linear order of overlapping preganglionic connections among the population of postsynaptic ganglion cells, which is thought to be based on chemospecificity. This pattern is similar to the juvenile linear order in muscle. On the other hand, which axon becomes the sole survivor at each muscle fiber is determined in a competition that may have more than one outcome. For example, in this work we noted that muscle fibers with two NMJs mostly ended up with different axons at each site, indicating that no single motor axon is the uniquely appropriate choice; nonetheless, all adult NMJs undergo synapse elimination until only one axon remains. Ample evidence supports the idea that activity differences among the innervating axons is an important determinant of the fate of axonal terminals (Busetto et al., 2000; Callaway et al., 1987; Favero et al., 2012, 2014; W. A. Harris, 1981; Personius & Balice-Gordon, 2001; Wyatt & Balice-Gordon, 2003). Our study adds a new argument for the role of activity in shaping the final pattern of connections that relates to the ultimate fate of the supernumerary NMJs on MEFs. We found that, as development proceeds, the relative (and absolute) distance between NMJs increases as the number of MEFs decreases. This means that NMJ elimination is less likely to occur if the junctions are far apart. Moreover, the last remnant of nearby neuromuscular junctions that are present in the second postnatal week always had the same axon innervating both sites. We show that there is a plausible mechanism to explain these results based on the conduction of action potentials from one neuromuscular junction to another and the prevention of this, if the activities of the two junctions is close enough in time for the action potentials to collide as they propagate (slowly) in the muscle fiber between the 2 endplates. Thus we propose that muscle fibers that retain 2 endplates do so because the activity at the two sites is so similar that they basically have no way of communicating due to action potential collision. Consistent with this view is the linear ordering of the axons that participate in the innervation of MEFs.

All of these results argue for a considerable amount of coordination in the connectivity of different axons. Unlike a strict molecular matching, the outcomes are graded and variable -- even when axons maintain synapses on the same muscle fiber they are usually not the exact same axon. Rather, a role for activity-based interaction between competing synaptic contacts seems to be regulating both synapse elimination within a neuromuscular junction and the perseverance or elimination of distant synaptic sites on MEFs. The fact that this coordination can impact synapses that are even 1 millimeter apart invites speculation about whether such mechanisms might coordinate inputs to neurons. For example, synapses on the basal and apical dendrites of pyramidal neurons in the cerebral cortex are separated by a long distance; however the intervening region of the cell is capable of generating back-propagating action potentials providing a system not entirely unlike a MEF. More generally we wonder if the recruitment order idea present in the neuromuscular system may provide insights into the ways other graded brain phenomena (e.g., the intensity of an emotional response) are organized. We suspect that the events that regulate which axons are eliminated and which strengthen on muscle fibers as they develop will have implications for many other parts of the nervous system.

## ACKNOWLEDGEMENTS

We would like to thank the Computational Connectomics Group (CCG) of Prof. Nir Shavit at MIT CSAIL for insightful discussions on input synchronization. We would also like to thank several individuals who greatly contributed to this work: Jincheng Tian for his enormous efforts to reconstruct the muscle fibers in the newborn sample, Vikram Norton for inspiring discussions and hard work on the reconstruction of several axonal arbors, Emma Yang for multifaceted contribution to the reconstruction of the newborn sample, Sophia Laskaris for insightful discussions, animal care and preparation of the machine learning datasets for the muscle fiber expansion and mitochondria classification, Tien Tran for axon reconstruction in the EM dataset, John Brady for assistance with manual reconstruction of axons from the EM datasets and inspiring discussions on multiple-endplate fibers early in this study, Marco Badwal for many discussions and his early contribution to the research, including reconstruction of the extra-muscular dataset, Brandon Drescher for help with the preparation of the myonuclei data for machine learning, Siyan Zhou for help with the initial analysis of the confocal image with the EM and reconstruction of Schwann cells in the nerve, and Xupeng Chen for helping with the reconstruction of part of the extra-muscular bundle and assistance with Schwann cell reconstruction.

## METHODS

### Sample acquisition and preparation of the newborn sample for ssSEM

Thy1-YFP16 mice (Feng et al., 2000) were bred and housed according to the guidelines of the Harvard Animal Care and Use Committee. In newborn pups (P0), deeply anesthetized with ketamine-xylazine, the interscutularis muscle was exposed and labeled with 5 µg/ml Alexa 647-conjugated alpha-bungarotoxin (ThermoFisher Scientific, USA) for 10 minutes at room temperature to stain acetylcholine receptors as a guide for neuromuscular junctions. Immediately after thoroughly rinsing the muscle with 0.1 M phosphate-buffed saline (pH=7.4, PBS, ThermoFisher Scientific, USA), the interscutularis muscle was immersed with a solution containing 2% glutaraldehyde (Glut., EMS, USA) and 2% paraformaldehyde (PFA, EMS, USA) in 0.1M sodium cacodylate buffer (pH=7.4) for 6 hours at 4°C. Then, under a Leica fluorescent dissecting scope (Leica, Germany), the interscutularis muscle was isolated and trimmed to only contain the entire endplate band and ∼1 mm of the posterior auricular nerve. After trimming, the sample was placed directly in the concave cavity of a glass slide and immersed in 0.1 M PBS to help keep the muscle from drying. A coverslip was placed over but did not touch the sample. Then, the sample was imaged using Zeiss LSM 710 confocal microscope equipped with Plan-Neuluar 10 × 0.3 NA objective. YFP and Alexa 647 were excited by 514 nm and 633 nm lasers respectively. The purpose of confocal imaging was to capture the nerve’s rough contour of innervation. It assisted further electron microscope image processing. We acquired tiled stacks of the entire endplate band scanning YFP and Alexa-647 simultaneously at a resolution of 0.923 µm, 0.923 µm and 3.017 µm for x, y and z dimensions respectively. After imaging, the sample was postfixed with 2% Glut and 2% PFA in sodium cacodylate buffer (0.1M, pH = 7.4) at 4°C for 12 hours, and processed for electron microscopy as described in previous work (Juan Carlos Tapia et al., 2012). Briefly, the sample was contrasted with ROTO (Reduced Osmium tetroxide-Thiocarbohydrazide-Osmium tetroxide; (Juan Carlos Tapia et al., 2012) and dehydrated with graded concentrations of ethanol (20%, 50%, 70%, 80%, 90%, and 100%; 15 min each), and propylene oxide (PO, 15 min twice). Finally, the sample was immersed in a mixture of 3:1 (30 min), 1:1 (1 h), 1:3 (3 h) of PO and Plain Resin (Nisshin EM, Tokyo, Japan), and infiltrated with pure Plain Resin for 12 hours. After replacing the resin at least four times, the sample was finally embedded in beam capsules and cured in an oven at 50°C for 24 h and 70°C for 4 days.

### Sample sectioning for ssSEM

The surface of the resin block was first trimmed to a rectangle, and the corners of the rectangle were then trimmed to form an arrow-point on each end of the now six-sided polygon block using a 3 mm ultratrim diamond knife (Diatome, USA) and a UC6 ultramicrotome (Leica, Germany). The fibers were spatially aligned with the length of the rectangle and sectioned perpendicular to the fiber direction in order to reduce sectioning compression. The 60 nm thick serial sections were cut with a 45° ultra diamond knife (Diatome, USA) and collected using the automated tape ultramicrotome (ATUM) system (Kasthuri et al., 2015). The sections were cut at a speed of 0.3 mm/s and collected onto carbon-coated Kapton tape. A total of 3,232 serial sections were collected. The first 2,315 sections contain muscle tissue and the nerve innervating the muscle. The following 917 sections and uncounted sections only contain the nerve and no muscle.

### Wafer fabrication and mapping

The tape holding the sections was cut into strips with a razor blade between collected sections and adhered with 25.4 mm wide double-sided conductive carbon adhesive tape (Ted Pella Inc., USA) onto 4 inch diameter circular wafers (University Wafers, USA). A total of 3,232 sections were distributed across 25 wafers. To enhance the signal from cell membranes, each wafer was first plasma-treated for 30 s (operating pressure of 1 × 10-7 mb, plasma current of 15 mA) to increase its hydrophilicity, immediately stained with 4% uranyl acetate for 3 min, rinsed with ddH2O for 30 s 10 times, air-dried, stained with 3% lead citrate (Leica UltroStain II) for 3 m, rinsed and air-dried as before, stored under vacuum. Each wafer was mounted on a metal wafer holder with fiduciaries to target high-resolution imaging by multi-beam scanning electron microscope (MultiSEM 505, Zeiss). A low-resolution (3.57 µm/pixel) optical image was acquired from each wafer mounted on the wafer holder, which identified the position of each section relative to the wafer holder fiduciaries (Hayworth et al., 2014). A six-sided polygonal region of interest (ROI) was defined and superimposed onto each section in the optical image of each wafer using the Zen software package (Zeiss Microscopy), to target high-resolution imaging using MultiSEM 505.

### ssSEM image acquisition

The first 18 wafers contain the interscutularis muscle and nerve fibers innervating the muscle, and the sections on these wafers were imaged using the multi-beam SEM, which employs 61 electron beams to scan 61 overlapping rectangular regions simultaneously (Eberle et al., 2015), producing 61 image tiles. These 61 tiles form one large hexagonal image known as a multi Field of View (mFoV). Once one mFoV has been acquired, the multi-beam SEM stage moves to an adjacent site to acquire another mFOV. All images were scanned at 4 nm/pixel resolution, with a tile overlap (within mFoV) of 0.5 µm and a between-mFoV overlap of 4%. Each ROI was imaged with landing energy of 1.5 kV with each scanning beam at 570 pA, and a dwell time of 3.2 µs/pixel. Brightness and contrast for each ROI were set to maximize the dynamic range of the images acquired, by maximizing the spread of the histogram of image grey levels without clipping its tails. Prior to imaging each ROI, the multiSEM was programmed to determine the optimal focus height and stigmation settings at ‘focus support points’ (FSPs) within the section. If this procedure failed at more than 25% of the FSPs, then the ROI was not acquired, and the procedure was restarted with new FSPs added; failed FSPs removed or moved to other locations. Once successful at 75% or more FSPs, Delaunay triangulation was used to interpolate a topological map of the ROI, to guide the autofocus of the multiSEM during the imaging of the ROI. The last 7 wafers with sections containing only nerve fibers were imaged using a single electron beam ZEISS Sigma. The image was scanned at 4 nm/pixel resolution, with landing energy of 1.7 kV at 1 nA, and a dwell time of 200 ns per pixel.

### Image Stitching and Alignment of the ssSEM sample

Stitching and alignment of the dataset presented some challenges due to the large spatial section area, the sparsity of features in many of the tiles and sections, the arc-like structure of the tissue, and the wrinkles that appear in random places throughout the sections. These issues were addressed by the techniques described below. The overall stitching and alignment was similar to Saalfeld et al. (Saalfeld et al., 2012), where each step was parallelized over image tiles and sections.

#### Stitching

To stitch the tiles of the sections, point correspondences were obtained between overlapping tiles, followed by an optimization process that estimates the tile position and rotation while minimizing the matching correspondences’ sum of squared errors. As a first approximation, the multiSEM stage position data acquired during the imaging phase was used to estimate the tile locations. The approximate locations were used to detect overlapping areas between pairs of tiles. Next, a CLAHE filter was used to increase the contrast between these areas (e.g., in low texture regions such as blood vessels), followed by ORB (Rublee et al., 2011) features computation in each of the areas. The features of the overlapping tiles were matched in order to find correspondences. To remove outliers of incorrect correspondences, the RANSAC algorithm (Fischler & Bolles, 1981) was used while optimizing for rigid transformations where the assumed rotation is less than 5 degrees. Finally, the optimization of the tiles’ rotation and position that minimizes the sum of squared roots of all the correspondences was done.

#### 3D Alignment

To align the stitched sections, coarse features were detected on each section, and matched between the sections, which were then used to guide the searches for patch matches. The matching was performed between all pairs of neighboring sections up to two sections apart. Coarse features were obtained from the original tiles, to avoid the rendering of the stitched sections. The features were detected using the OpenCV SimpleBlobDetector (“OpenCV: cv::SimpleBlobDetector Class Reference” n.d.). The detector searches for circular dark blobs, corresponding to lipids in the tissue, which appear in the same spatial location in 4-7 consecutive adjacent sections. SIFT descriptor (Lowe, 1999) was used to describe the features, and after matching the features of a pair of sections, RANSAC was used to filter out wrong matches and to find an affine transformation between the sections. Next, fine-grained patch matching at half resolution was performed. To this end, a triangular mesh was laid on the sections, 1.6×1.6 µm^2^ area around each mesh vertex was cropped, followed by cross-correlation search against a transformed cropped area of the neighboring section of size 4×4 µ m^2^. The valid cross-correlation matches were then used as an input to a 3D elastic optimization algorithm based on prior work (Saalfeld et al. 2012), which minimizes the cross layers distance of the matches while preserving the 2D structure of the sections.

### Semantic and instance segmentation of the Newborn ssSEM sample

#### Endplate identification

Three annotators exhaustively searched for endplates by scrolling through the volume in VAST at a low mip-level, identifying axon bundles and presynaptic vesicle-filled swellings. Three independent strategies were adopted for endplate identification: one annotator followed fibers along all their length, another annotator searched for axon terminals and a third annotator extracted candidate locations based on locations indicated by the confocal stack of the tissue stained with alpha-bungarotoxin to identify AChR sites.

#### Muscle fiber reconstruction

An expert annotator enumerated and seeded all myofibers that intersected the right side of the volume. This approach required to scroll through the z-axis, identify the rightmost extreme of a myofiber, name it and volumetrically segment part of the fiber to assist visual search of other myofibers. This resulted in the identification of 208 muscle fibers which were skeletonized by a second annotator. Subsequently, the remaining 32 fibers were identified by backtracking fibers after all the initially seeded fibers were completely skeletonized in the volume. This backtracking was initiated both from identified endplates that were not associated with a skeletonized fiber and by a third annotator by exhaustive search for missing fibers in the horizontal middle plane of the dataset. All of the 32 fibers that were identified in the second step were: secondary fibers that did not reach the right side of the dataset, a small number of damaged fibers (∼5) or fibers at the periphery of the muscle that exited the volume at a slightly more medial point relative to the initial saturated box. An additional set of cellular structures with a topology similar to myofibers was identified as being composed of myoblasts, given the absence of sarcomere structure, their shorter extent and irregular sarcolemma. Interestingly, all the secondary fibers possessed an endplate close to the location of the endplate of the primary fibers to which they were morphologically associated (Fig. S12) as reported by (Duxson et al., 1989).

#### Volumetric instance segmentation of muscle fibers

To obtain a volumetric representation of the muscle fibers we trained a neural network to detect the sarcolemma (membrane of the muscle cells), sarcoplasm (the cytoplasm of the muscle cells), and voxels that did not correspond to any of the two categories. We then applied the neural network to the entire space using mEMbrain (Pavarino, E., Berger, D. R., Morozova, O., Drescher, B., Bidel F., Kang K., Lichtman J. W., Meirovitch Y., 2019) and obtained for each voxel affine scores for the three categories. We then used the human annotation as seeds for a 3D watershed algorithm. Pre-processing consisted of Gaussian smoothing of the probability maps. Post-processing of the 3D-expanded objects included removal of object voxels that were far from initially annotated regions, majority voting across sections, spherical filling around annotated regions that were not expanded.

#### Myonuclei Reconstruction

The polynuclear nature of muscle fiber cells is thought to control transcriptional activity in segmented domains of cytoplasm surrounding such nuclei. This myonuclear domain varies between fiber types (Qaisar & Larsson, 2014) and is inversely associated with the fiber’s oxidative capacity (Qaisar & Larsson, 2014; Van der Meer et al., 2011). Additionally, in vertebrate skeletal muscle myonuclei cluster at the muscle fiber endplates, forming aggregates of synaptic nuclei with distinctive function and morphology (Grady et al., 2005). Hence, comparing the myonuclear domains of multiple endplate fibers (MEFs) and single endplate fibers (SEFs) could provide us with insights about key differences that may exist between these types of fibers. To approximate myonuclear domain, i.e. (Volume of Fiber)/(Number of Nuclei in Fiber) in the newborn sample, we approximated it by Constant*(Fiber CSA) / (Number of Nuclei in Fiber) under the assumption that fibers maintain a relatively fixed CSA and that all fibers have similar lengths. We reconstructed all the synaptic and nonsynaptic myonuclei of 52 (25 MEFs and 27 SEFs) randomly selected muscle fibers in the newborn EM sample. In total we reconstructed 930 nuclei. To do so, one annotator placed skeleton nodes in each of the nuclei. We then trained a neural network using the software mEMbrain to classify pixels based on three classes: nucleoplasm, nuclear membrane, and pixels that do not belong to the two first categories. The output predictions of such a network were used by a second annotator, who expanded each seeded nucleus leveraging VAST’s flood-filling tool and using the classified pixels as constraints. In particular, this was achieved by clicking on the nucleoplasm while the computed nucleoplasm probabilities are set as the source layer of the flood-filling algorithm in VAST. This required adapting the threshold for the filling operations based on the quality of the expansion, in order to avoid a too limited expansion (i.e., not reaching the nuclear membrane) or an expansion that would spill through the nuclear membrane to the sarcoplasm. At times this reconstruction method required human correction e.g. fixing the expansion by erasing where it had overflowed the nuclear membranes. Nevertheless, this semi-automatic procedure was several fold faster compared to the fully manual annotation of these objects. Finally, the nuclei were visualized in 3D and further analyzed.

#### Axon instance segmentation

##### Nerve

All axons were enumerated and manually skeletonized from wafer 12 to wafer 25 for over 1827 sections. This analysis identified a single fascicle containing nerve cells and Schwann cells, surrounded by perineurium tissue. Low-magnification inspection of the EM suggested no other fascicles projecting to the interscutularis muscle. To get a better idea of the number of distinct axons in this nerve, we identified and analyzed several cross sections of that nerve to the muscle for about another 1000 sections distal from the muscle (close to 200 um distal to the muscle fibers). The number of individual axonal cross sections in these slices matched the number of axons identified in the fully reconstructed nerve segment, i.e. 21 axons. The manually skeletonized axons were automatically expanded to a faithful volumetric 3D reconstruction using mEMbrain (Pavarino, E., Berger, D. R., Morozova, O., Drescher, B., Bidel F., Kang K., Lichtman J. W., Meirovitch Y., 2019) and adopting a similar approach to the fiber reconstruction, as explained above.

##### Axon bundles

The 21 axons identified in the nerve reconstruction were traced throughout the muscular tissue by seven annotators (three experts and four extensively trained undergraduate students). The estimated total number of annotation hours exceeded 3500 human annotation hours. The first task was to skeletonize the axonal bundles as deeply as possible iteratively across all possible bifurcations using the annotation software VAST. Once the procedure was saturated, three annotators commenced tracing from identified endplates, connecting anonymous axon segments to identified fascicles, until all axonal branches were either associated with a synapse (the vast majority of cases) or rarely reached specialized zones containing disorganized axonal swellings surrounded by intricate glial processes close to the sarcolemma of a muscle fiber. We were unable to exclusively determine the nature of these regions with the hypothesis that they are related to sites of muscle innervation from the embryonic stage which we speculated might be associated with early stage axonal retraction from neuromuscular junctions.

### Sample preparation of brainbow mice

#### Transgenic brainbow animals

To label motor neurons in multiple colors, we initially tried ‘first generation’ multicolor transgenic strategies that relied on cytoplasmic expression of fluorescent protein (Livet et al., 2007). These strategies fell short because of the limited number of distinguishable colors or low expression levels early in postnatal life. To have early onset of bright labeling in many colors, we developed new lines of Brainbow transgenic mice in which all fluorescent proteins were membrane tethered (Fig. S13). Because changes in axon caliber have a greater effect on volume than on surface area, we reasoned that the membrane label might be brighter and more uniform than the cytoplasmic label for fine neural structures at any given expression level. To create the multi-color membrane label, we generated Brainbow mice in which three fluorescent proteins (eCFP, eYFP, mCherry) were directed into the plasma membrane by an N-terminal palmitoylation tag (Kay et al., 2004). We crossed these mice to Hb9^cre^ transgenic mice, which express Cre recombinase in motor neurons postmitotically (Arber et al, 1999). *In vivo* expression of the fluorescent proteins was indeed restricted to the plasma membrane. However, the fluorescent protein was localized primarily to the axon and was almost completely absent from the soma and dendrites. This compartmentalized expression likely contributed to increased brightness of motor axons.

Thy1-Membrane-Brainbow animals were generated as described (Tsuriel et al., 2015) and crossed to Hb9-Cre animals obtained from Jackson labs (JAX stock 006600).

#### Histology (brainbow)

Mice were deeply anesthetized with sodium pentobarbital and perfused transcardially with ice-cold 4% paraformaldehyde (Electron Microscopy Sciences) in PBS. Muscles were dissected and post-fixed in 4% paraformaldehyde for 30 min. Muscles were rinsed with 0.1 M glycine in PBS for 10 min, then 5 min in PBS and incubated for 30 min in 5 µg/µl Alexa 647-conjugated bungarotoxin (Thermo Fisher Scientific) dissolved in 1% bovine serum albumin (Sigma-Aldrich) in PBS. Muscles were rinsed again in PBS for 15 min. All post-fixation was done at 4oC, rocking in the dark. Finally, muscles were mounted on slides in Vectashield H-1000 fluorescent mounting media (Vector Laboratories), flattened with magnets overnight at -20 oC, then sealed with nail polish (Electron Microscopy Sciences), and stored in a freezer (−20oC).

To collect a sample where all axons had unique colors, we screened hundreds of animals, and analyzed muscles where the number of color redundancies among axons at the muscle entry site was limited to one or two. We always found more axon collaterals than colors at the muscle entry site. It is likely that some color redundancies were due to branching of the axon in the distal nerve (see Lu et al, 2008). In seven muscle samples the collaterals with identical colors 1) never co-innervated the same neuromuscular junction, 2) did not have motor unit sizes twice the average size of others, and 3) displayed similar skewing in partner preferences (see below). For these reasons, these identically colored axons were likely axon collaterals rather than statistical color redundancies. Thus, these muscles contained color segmented axons that could be analyzed as complete connectomes (Fig. S14): four omohyoid muscles (P6, six axons; P7, nine axons; P7, six axons; P8, five axons) and three forepaw lumbrical muscles (P8, six axons; P9 six axons; P9, six axons).

#### Imaging of brainbow samples

Images were taken using the Multi-Area Time Lapse function of the Olympus FV1000 confocal microscope (Olympus Scientific Solutions Americas Corp., Waltham MA) with 3% overlap between adjoining tiles using either a 60x 1.4 NA PlanApo or 1.35 NA UPlanSApo oil immersion objective lens. The 1024×1024 and 800×800 frame size images were zoomed at 1.9 and 2.4, respectively, and stepped in Z by 0.37 µm. The 800×800 frame size was used when the image stack was greater than 121 sections due to memory limitations of the computer operating system. Samples were excited in two sequential groups to reduce spectral bleed-through: first with the 440- and 633-nm laser lines and second with the 515- and 561-nm lasers. Emission filters for four spectral channels were set as follows: Cerulean: 480/15 nm, EYFP: 535/15 nm, dTomato and mCherry: 600/25 nm, Alexa 647: long pass 650 nm. Typically the images were collected at 2 µs/pixel with either no averaging or kalman = 2.

#### Identification of presynaptic innervation in the confocal brainbow stacks

To assess the identity of motor axons at each junction, each image stack was individually evaluated using the ‘section view’ panel in the Imaris software. This panel displays a maximum intensity projection of a re-sizable subvolume of the data stack and allows real-time, user-specified gamma and threshold adjustment. By re-sizing and scrolling a small sub-volume maximum intensity projection of the data stack through the Z-dimensions, the color of each non-overlapping axonal terminal at or near the synapse could be compared to other colors present in the surrounding volume. When no non-overlapping region was available or the structure size was too small to accurately judge the color of the process, the neurite was followed back along its path to see more samples of its color and branches to which it connected. In cases where tracing was obscured by other axons of similar color or low contrast, the identity of the synapse was left “ambiguous”. This occurred in less than 5% of the total junctions.

### Sample preparation of YFP16 mice

Neonatal (Postnatal days 1, 3, 5, 10, 12) and Young adult *thy-1*-YFP-16 transgenic mice (P30 and P60) were anesthetized via IP (intraperitoneal injection) of 0.1ml / 20g ketamine-xylazine (Ketaset, Fort Dodge Animal Health). Following deep anesthesia, animals were transcardially perfused with 4% p-formaldehyde (PFA; Electron Microscopy Sciences, USA) in 0.1 M phosphate-buffered saline pH 7.4 (PBS; Sigma-Aldrich, USA). The interscutularis muscle, along with a ∼1 mm segment of the posterior auricular nerve was dissected out, and postfixed in the fixative solution (4% PFA in 0.1M PBS, 1h). After several rinses (in PBS), samples were incubated with alpha-bungarotoxin conjugated with Alexa594 (α-btx594, 4h; Thermo Fisher Scientific) to stain the acetylcholine receptors (AChRs). Then, after removing the excess of α-btx594 with PBS (30 min, 3x), samples were mounted on slides with Vectashield mounting medium (H-1000, Vector Laboratories). In some samples, Wheat Germ Agglutinin (WGA) conjugated with Alexa647 (WGA647; ThermoFisher) was used to label muscle fibers contours, and the overall muscle vasculature. All samples were imaged using either Olympus FV1000 (Olympus Scientific Solutions Americas Corp., Waltham MA) or Zeiss LSM-710 (Carl Zeiss Microscopy, LLC, White Plains NY) confocal microscopes as indicated below.

### Confocal Imaging of the developmental and adult samples

We imaged whole-mounted neonatal and adult mouse muscle using Zeiss LSM-710 (Carl Zeiss Microscopy, LLC, White Plains NY) and Olympus FV1000 (Olympus Scientific Solutions Americas Corp., Waltham MA) laser scanning confocal microscope systems that were equipped with a motorized stage, high numerical aperture oil-immersion objectives (Zeiss Plan-APOCHROMAT 63x/1.4; Olympus UPlanFL N 40x/1.3 and Olympus UPlanSApo N 60x/1.35), and spectral channels to detect the fluorescence emission. The Zeiss and Olympus systems were running ZEN 2010 and FV10-ASW 4.1, respectively, and had optional software modules to manage acquisition of tiled image stacks (Zeiss: LSM StitchArt; Olympus: Multiple Area Time Lapse). We used a multi-band primary dichroic filter to reflect the laser lines needed to excite YFP, Alexa 594 and Alexa 647 (514.5 nm, 561 nm and 633 nm, respectively). For each muscle (p1 to p60), we acquired tiled stacks of the entire endplate band, simultaneously scanning YFP and Alexa-647 and sequentially scanning Alexa-594. The voxel size was isotropic in X, Y and Z (e.g., p1: 0.31 µm, p10: 0.42 µm and p30: 0.83 µm) to facilitate orthogonal- and off-axis slicing in subsequent analysis to count/track the mapping of endplates to muscle fibers. The Zeiss tiled datasets were montaged using the stitch function of the LSM StitchArt module. The Olympus tiled datasets were montaged in Fiji (Schindelin et al., 2012) using the Grid/Collection stitching plugin (Preibisch et al., 2009). For p60 interscutularis muscle, we also imaged YFP at very high resolution on the Olympus system using the 60x objective (0.138 µm per pixel in XY and 0.2 µm in Z). The high-resolution single-channel image volume allowed tracing of the full arbor of each YFP-filled axon of the motor input (i.e., the construction of the muscle connectome when combined with the corresponding lower-resolution multi-channel image volume).

### Reconstruction of muscle fibers and axon arbors and endplates from the confocal stacks

#### Muscle fibers, endplates, and multiple-endplate fibers

All confocal image stacks were manually annotated by two expert annotators. For the P0, P3, P5, P10, P12, P30, and P60 samples, muscle fibers were enumerated and saturated in two planes perpendicular to the muscle fiber axis at about one and two-thirds portions of the volume across the longitudinal axis of the muscle. In all samples, the respective two counts yielded the same number of muscle fibers plus or minus ∼5 fibers. In P0-P5 the two estimates were averaged and used as the estimate of the number of muscle fibers in each volume. In P10, P12, P30, and P60 the muscle fibers were manually reconstructed (or in one sample automatically reconstructed using mEMbrain (Pavarino et al., 2019) and manually proofread). In all samples, the number of endplates was defined by firstly identifying potential endplates from the 3D rendering of the red (α-btx) and green (YFP16) channels of the volume in VAST, by identifying volumetric structures of a stereotypic endplate shape. At earlier ages (P0-P5), all high-energy regions with an intense red-channel signal (α-btx) were added to the list of candidate endplates. During this process, in conjunction with careful analysis of each case, we realized that all of the identified endplates are in the vicinity of axonal bundles. We, therefore, were able to identify and skip noisy aggregation of red-channel high intensity pixels that were located far from the axonal tree. Each of the putable endplates was also analyzed by traversing the 2D image stack on transverse orientation and verifying the convexity of the perimeter of the endplate (α-btx) around the identified muscle fibers (blue-channel).

The difference between the number of identified endplates and the number of identified muscle fibers provided us with an estimate of the number of multiple-endplate fibers and their proportion in each sample. In P10-P60 we also directly classified each of the muscle fibers as multiple-endplate or single-endplate fibers by swiftly animating the 2D stack across the longitudinal axis in VAST. Both measurements gave us the same number in the vast majority of cases; when small discrepancy occurred (never more than ∼5 fibers), the procedure was repeated until the source of discrepancy was identified (either an endplate was missed in the initial search for endplates from the 3D rendering, or a muscle fiber was wrongly classified during the subsequent procedure based on 2D search).

#### Axon tracing

Tracing the axonal arbors of two adult interscutularis muscles (P60) was accomplished in VAST (Berger et al., 2018) using manual segmentation by one expert annotator aided by a second annotator. Each annotator focused on a different part of the dataset. This was made possible due to the very high resolution YFP-based confocal images at 138 nm/pixel. The tracing strategy that we found most convenient was to trace axons in their transverse sections which in most cases required using an orthogonal plane annotation plane in VAST. When tracing became difficult (rarely in these high-quality stacks), to avoid ambiguity, the two annotators raised concerns and attempted to reconcile contradicting opinions. Several regions required postponing the decision for a difficult axonal tracing until the companion axons, at the vicinity of the ubiquitous axon reconstruction, were reconstructed. In all cases, this led to an agreeable solution.

### Statistical Tools and Analysis

#### Linear order of axons

Whether the innervation of muscle fibers by motor neurons is initially unspecific (random in nature) or specific (each subpopulation of axons has a greater tendency to innervate a particular sub-population of fibers) has been a topic of a number of theoretical studies (Willshaw, 1981). The main tenet has been that while the outcome of synapse elimination at each NMJ depends on a variety of factors, the cohort of axons innervating each junction likely reflects their developmental history. For example, the opportunity to innervate a junction by a specific axon may be related to the limited axonal topography at a developmental time, and possibly also to the status of innervation of nearby junctions. Wilshaw specifically hypothesized that the motor system benefits from a random innervation pattern that is purposefully excessive at birth, only to allow each junction a sufficiently high probability to maintain at least one input (Willshaw, 1981).

We assessed the level of randomness in the innervation pattern by computing for each pair of axons the probability to have a certain number of shared muscle fiber targets, under the null hypothesis that axons are equally likely to dispense their terminals at all junctions independent of each other. Indeed, in the newborn interscutularis muscle all axons, except one, are arborized at birth across the entire longitudinal axis of the muscle. All axons also innervated endplates in more or less equal numbers at the muscle regions distal and proximal to the entry point to the muscle tissue. Nonetheless, it is clear that even if endplates are innervated at equal likelihoods, the probability to innervate one endplate is not independent of the probability to innervate other endplates (the mutual information is not zero). Hence, we devised a tool to assess the global innervation pattern based on the deviations of pairs of axons from the null hypothesis that for an axon with K muscle fiber targets, all subsets of K targets are equally likely to occur. We defined the link between two axons based on how excessive the number of targets shared between them is compared to a random innervation model. Formally, we considered for two axons with N1 and N2 targets the probability to observe exactly S shared targets assuming there exist M equally likely innervation sites.

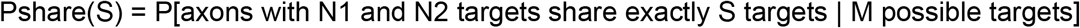

We then summed the probabilities to observe the actual number of shared targets or beyond.

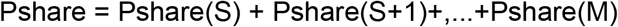

This yielded a network of probabilities among all pairs of motor axons. For example, axons 17, 19, and 21 assumed a sub-network as depicted in Figure S18. Of the 240 muscle fiber targets in the muscle, axon 17 innervated 93 fibers (innervation ratio ∼39%) and axon 21 innervated 90 fibers (innervation ratio ∼38%). If their muscle fiber targets are innervated with equal likelihood and independently of each other, we would expect that one axon will innervate the targets of the second axon with the same ratio of innervation as it innervates the entire muscle. However, we found that the number of shared targets, 54, is ∼58% and ∼60% of the number of muscle fiber targets innervated by the two axons, respectively. The probability to observe this number of shared targets, or more, for motor unit sizes 93 and 90, and for a total of 240 possible targets, assuming independence of innervation, is about 387*10^−10^ (Wang et al., 2015). As we do not assume that the probabilities to innervate two junctions are independent of each other, we will use the above probability Pshare as a score for the tendency of two axons to innervate many targets. This can be seen as a way to deal with the number of shared muscle fiber targets of two axons with a normalization for the prior (fixed from the data) for the number of targets of each axon, and the total number of available targets. Similarly, we also compute the probability Pavoid by considering the number of targets that are not innervated by each axon and the number of targets that are jointly not innervated by the two axons (the complement of the overlap and motor unit sizes).

We were interested to see if the tendency of axons to innervate the same targets or avoid innervating the same targets is part of a global regularity of axonal networks. To this end, we defined the Sharing-network as a weighted graph of N axonal nodes, and edge weights between all pairs of axonal nodes defined by the negative log probabilities to observe the data (i.e., -log(Pshae), (Wang et al., 2015)). The larger the weight, the less likely it was to observe the number of shared targets (or more) by chance. Similarly, we build a network Avoidance-network to assess the tendency of axons to avoid innervating the same targets. When depicting these networks we observed an unexpected graph regularity which, except for one extremely sparse connectome, appeared in all the developmental ages for which complete connectomes were obtained in our data (P0 n=1; P6-P9, n=7; and P60, n=2). The regularity was shown as a tendency of axons to regularly innervate the same targets forming a special form of axonal neighborhood, with a linear appearance (axon_1_, axon_2_, …,axon_K_) or in one of the adults even simpler many axons clustered together repeatedly innervating the same set of multiple-endplate fibers. To capture this regularity and assess its statistical significance, we measured the wellness of embedding the axonal network on a line such that nearby axons tend to share the same targets (the weight from the Sharing-network) and a small tendency to avoid targets (the weight from the Avoidance-network). To do so, we first computed a 2-dimensional embedding with a spring-force algorithm that attempts to position the axons in the plane while respecting their pairwise innervation pattern (Force algorithm). We then measured the linearity of this embedding using the ratio of variances along the main axes of the embedding, formally the ratio of the first and second eigenvectors of the covariance matrices of the optimized positions.

#### Comparison of the empirical connectomes to graph distributions

If not stated explicitly otherwise, all the statistical assessments were carried out by comparing the developmental connectomes to theoretical graph distributions. This allowed us to derive the probability to observe the data (or more severe case) under the (null) assumption that developmental connectomes share properties with random graphs. This approach was undertaken when a permutation test that considers alternative assignments for a specific connectome (connectivity matrix) would not have been fruitful in assessing the statistics. We considered three main graph distributions.

Most commonly we used the configuration model where graphs are derived by considering a fixed number of nodes (usually for axons and muscle fibers) and edges represent the connections between them. In the asymptotic approach, to draw a graph instance from a distribution, we toss a coin with a positive probability that is proportional to the fixed degrees of the node in the data (Britton et al., 2006). In this approach, we consider a family of connectomes whose average node degree (i.e., number of connections incident to a node) is identical to the node degree in the data (e.g. number of synapses in a connectome or the number of shared targets between two axons in a co-innervation network). This approach is suitable when our biological assumption is that the number of edges (most often synaptic connections) may assume a considerable variability if many biological samples were derived. This was the case in the newborn connectome where also within the sample some muscle fibers were innervated by three axons and some by fifteen axons, and some axons innervate 13 muscle fibers and some innervate 120 muscle fibers. We do not wish to restrict the analysis to connectomes that identically follow the motor unit sizes and convergence numbers as in the data. This approach was used to produce random connectomes on which the linear order property was computed.

For the developmental ages and for the adult, allowing the variability in node degrees of the above will consider networks that are biologically irrelevant. For example, having a distribution that in average produces node degrees of one and two edges per node in the adult connectomes will allow for specific instances random connectomes with 3-4 inputs (with non negligible probability) - we however prefer to consider cases that are more consistent with what we expect may happen across samples. Hence, for these ages we used a shuffling approach that produces connectomes with exactly the same motor unit sizes and convergent numbers on fibers (or on endplates) as in the data. Also on these graphs we computed the relevant graph properties and derived the estimated p values (whose confidence intervals are known as well), including the linear order of the induced network of co-innervating axons.

In some cases where the developmental connectome assumed a special property we were interested to test some graph properties against random connectomes that have that special property and against such graphs that do not have that property (for example the connectome of the adult Muscle B which was highly central). In such cases we computed both probabilities (against different hypotheses) and described the outcomes under the different assumptions. For example when we assessed the motor unit sizes occurring on the adult muscle B we first discovered that the motor unit sizes of axons innervating multiple-endplate fibers in that sample is larger than what occurs in random graphs of exactly the same configuration (node degrees). However, when we parsed the distribution according to the sub-distribution of networks with centrality as in muscle B that phenomenon was no longer unlikely - hence we cannot exclude the possibility that centrality in that muscle is the cause for the appearance of large motor units innervating the multiple-endplate fibers, and in fact such large motor unit sizes can be explained from the centrality alone. In other cases the specific permutation tests or parametric tests are explained in the text next to the result.

#### Modeling AP collision as the mechanism to maintain multiple endplates on the same muscle fiber

We consider a model of AP collision with the following assumptions. 1) When an axonal AP arrives at the presynaptic terminal, it reliably triggers an AP in the muscle fiber, with a synaptic delay *T_d_* between the axonal AP arrival and the onset of the muscular AP. 2) The synaptic delay *T_d_*is identically and independently distributed (i.i.d.) at the two endplates on the MEF, following a Gaussian distribution *N*(µ, σ^2^), where µ is the mean synaptic delay and σ is its standard deviation. 3) The muscular AP propagates with a constant speed *v*_*m*_ irrespective of the direction of propagation. We first consider the case in which the two endplates are innervated by two branches of the same axon. Let the distance from the branching point to the NMJs be *l*_*1*_ and *l*_2_, respectively. Let the distance between the two NMJs be *l*_*m*_, so the time for the muscular AP to propagate from one NMJ to the other is given by 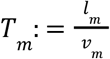. We define the moment when the axonal AP passes the branching point as time *t =* 0. AP collision occurs if, before the muscular AP elicited at NMJ #1 reaches NMJ #2, another muscular AP has been elicited at NMJ #2 but has not arrived at NMJ #1, and vice versa. The time at which the axonal AP elicits a muscular AP at NMJ #1 is 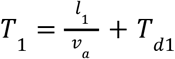. Similarly, the time at which the same axonal AP elicits a muscular AP at NMJ #2 is 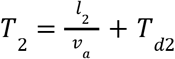. The time difference must be smaller than the time it takes for the muscular AP to propagate from one NMJ to the other:

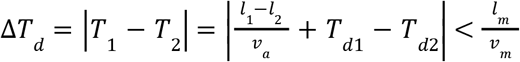

We may define 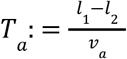 as the delay between axonal AP arrival time due to disparate path length. As *T*_*d1*_ and *T*_*d*2_ are i.i.d. Gaussian random variables, their difference *T*_*d*_ : *= T*_*d*1_ – *T*_*d*2_ is also a Gaussian random variable with mean µ_*d*_ *=* 0 and 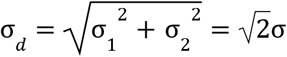. This gives the probability of collision:

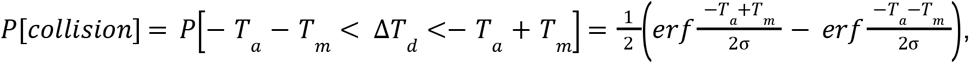

where erf is the error function. This equation also holds for the case in which two different axons innervate the same MEF at two junctions. In this case, *T*_*a*_ refers to the relative latency between the APs in the two axons.

## SUPPLEMENTARY INFORMATION

### EM correlates of neural degeneration

One of the advantages of carrying out such an exhaustive anatomical study with nanometer resolution is the ability of charting neural connections that might not be functionally active.

Throughout our p0 EM dataset, we observed signs of retraction. This is in line with findings from (Juan C. Tapia et al., 2012) who showed that the peak of motor unit size is just before birth (E18). We observed different levels of retraction: on one hand, we saw retraction of single branches of specific motor neurons (Fig. S16), while on the other hand, we saw retraction of entire sub-bundles of the neural circuit (Fig. S17). In the first case, specific anatomical features typically associated with retraction were circumscribed to local branches of specific axons. In particular, there were many axonal bulbs throughout the dataset, which were either found at the synapse or branched off the main axonal path (Fig. S16a). It is worth noting that these bulbs were devoid of synaptic vesicles, however they likely harbored axoplasm, traces of membrane recycling, or a higher concentration of lesser identified black dashes, which we hypothesized could have been either endoplasmic reticulum, or signs of cytoskeleton dismantling (see (Brill et al., 2016)). Another axon-specific sign of retraction was axonal local portions filled with vesicles (Fig. S16c), in alignment with previous studies (Bishop et al., 2004; Riley, 1981). Importantly, all axons surrounding the vesicle-baring axon did not present vesicles in the same proportion (or at all).

As reported above, one interesting phenomenon observed in our p0 dataset was signs of retraction – and perhaps neural degeneration – encompassing entire neural sub-bundles. From a morphological perspective these sub-bundles presented swellings in all the axons. It was not uncommon to see these axonal swellings filled with the aforementioned black dashes, or with signs of membrane recycling (Fig. S17b,c,d). The reason why we believe that these branches were retracting is that no axon of these sub-bundles was forming a synapse with muscle fibers. Furthermore, there was no evidence of synaptic vesicles within the motor neurons, suggesting that these terminals were no longer active, and hence a likely hypothesis is that they were retracting. These sub-bundles did not form synaptic contact with muscle fibers, but rather they terminated their journey in what we named “graveyards”, i.e. debris made up of Schwann cell and axonal material (Fig. S17a,b). Taking a closer look at samples of graveyards, we observed just how many of these shreds of axons contained a relatively high concentration of organelles, such as mitochondria and even vesicles. Moreover, the black dashes became prominent and more concentrated (Fig. S17b). While locating these graveyards, it was worth noticing that many were positioned in proximity of muscle fibers. Some of these fibers presented a higher concentration of mitochondria and lipid droplets, perhaps the heritage of a recently active postsynaptic site (Fig. S17b), while other fibers did not show any correlates of active zones (Fig. S17a).

Lastly, an intriguing anatomical correlate of our hypothesized retraction both at the axonal and at the sub-bundle level was the presence of putative degenerating synapses, devoid of synaptic vesicles. In some cases, these were present at the NMJ site (axon-specific) (Fig. S16b), while in other cases they were seen interfacing with non-active areas of muscle fibers (Fig. S17e,f).

## EXTENDED DATA

**Figure S1.**
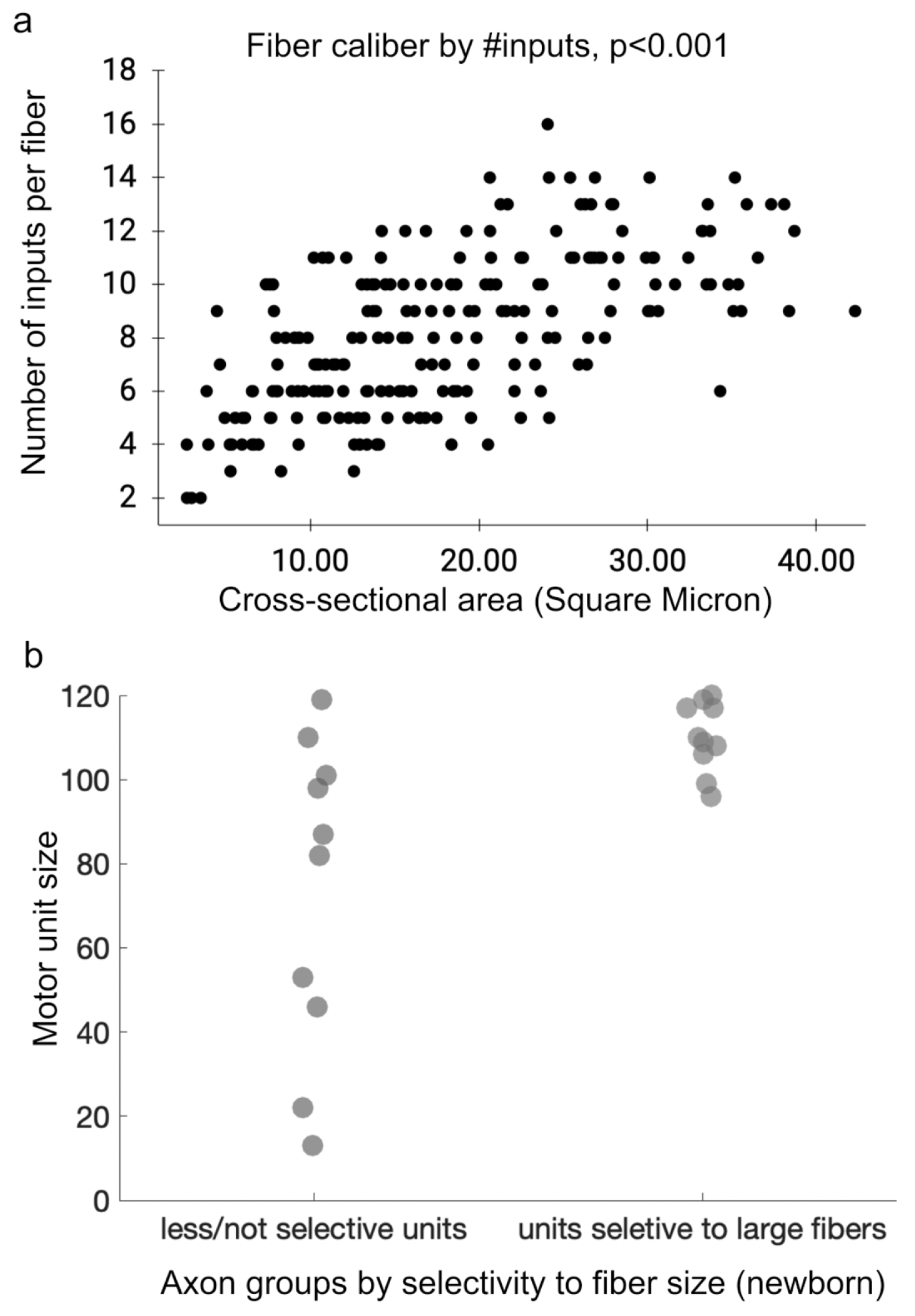
Association between muscle fiber size, number of inputs and motor unit size in the newborn. **a**. A correlation between cross-sectional area of muscle fibers and the number of distinct axons innervating them. **b**. Half of the motor units (n=10) were selectively innervating large muscle fibers (p<0.01); none of the small motor units was selective to large fibers and ∼half of the non-selective units were large motor units.

**Figure S2.**
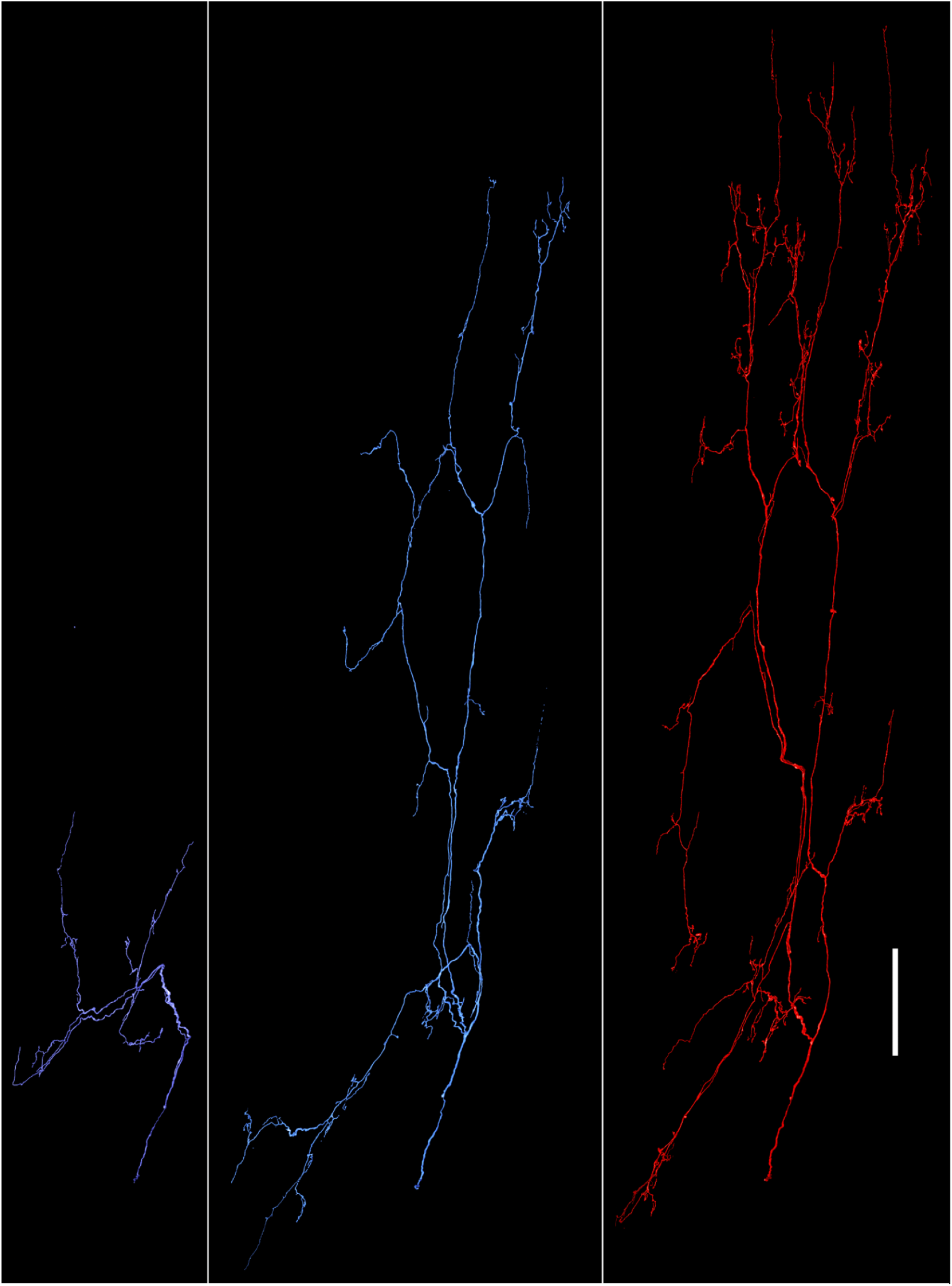
Varying sizes of axonal arbors in the newborn. Placeholder - we can add the endplates, and also in each rendering can add other arbors in transparent gray. The middle motor unit is contacting half as many fibers as the unit on the right although both axons are extensively extended across the innervation band. The middle axon proliferates less extensively from the main axonal branches.

**Figure S3.**
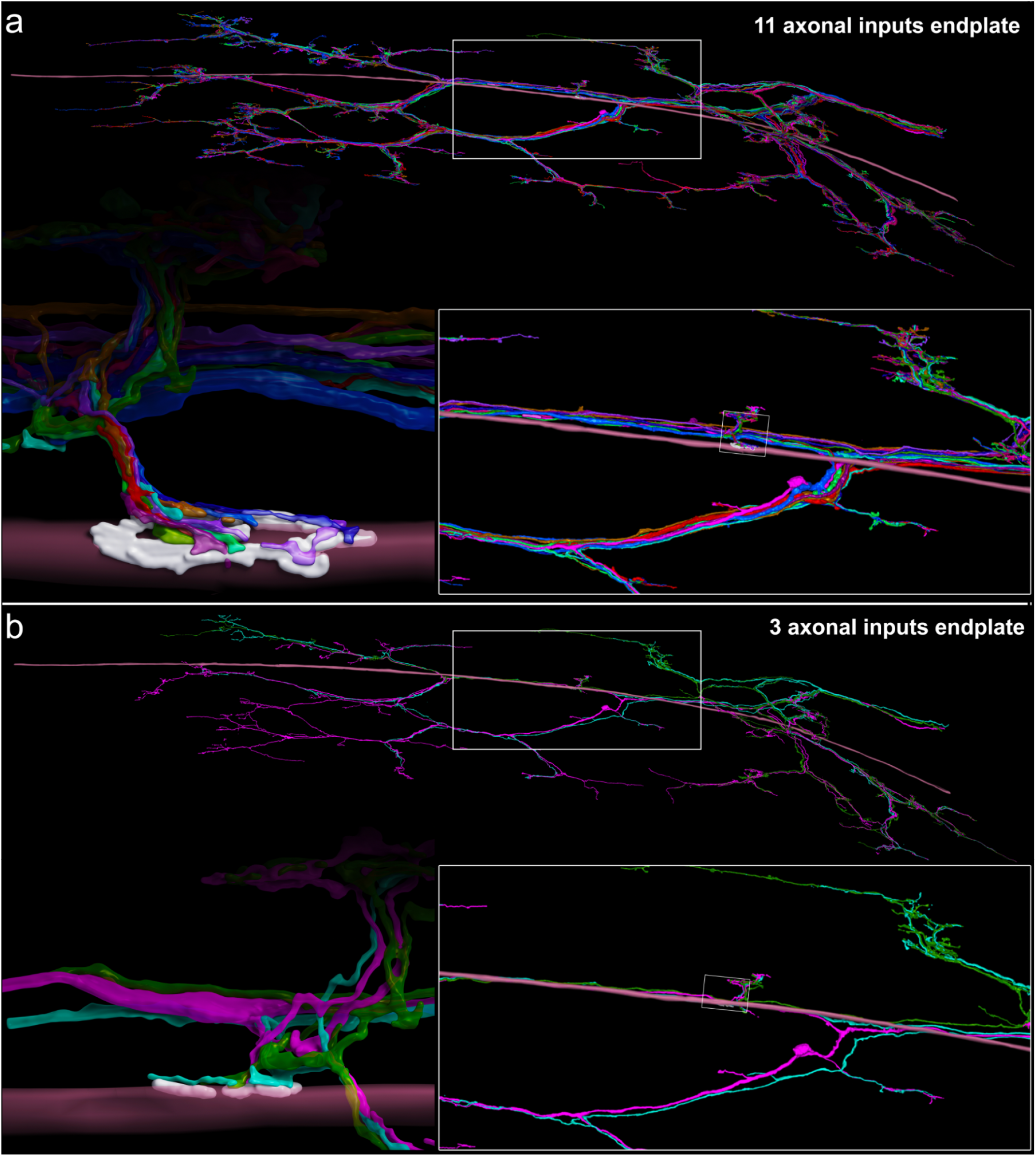
Large and small motor endplates in the newborn dataset. A rendering of an endplate (white) along their inputs are shown: **a**, 11-inputs endplate and **b**, 3-inputs endplate. The innervated muscle fiber and innervating axons are shown using random colors. The zoomed-up rendering of the endplates are shown with the same scale.

**Figure S4.**
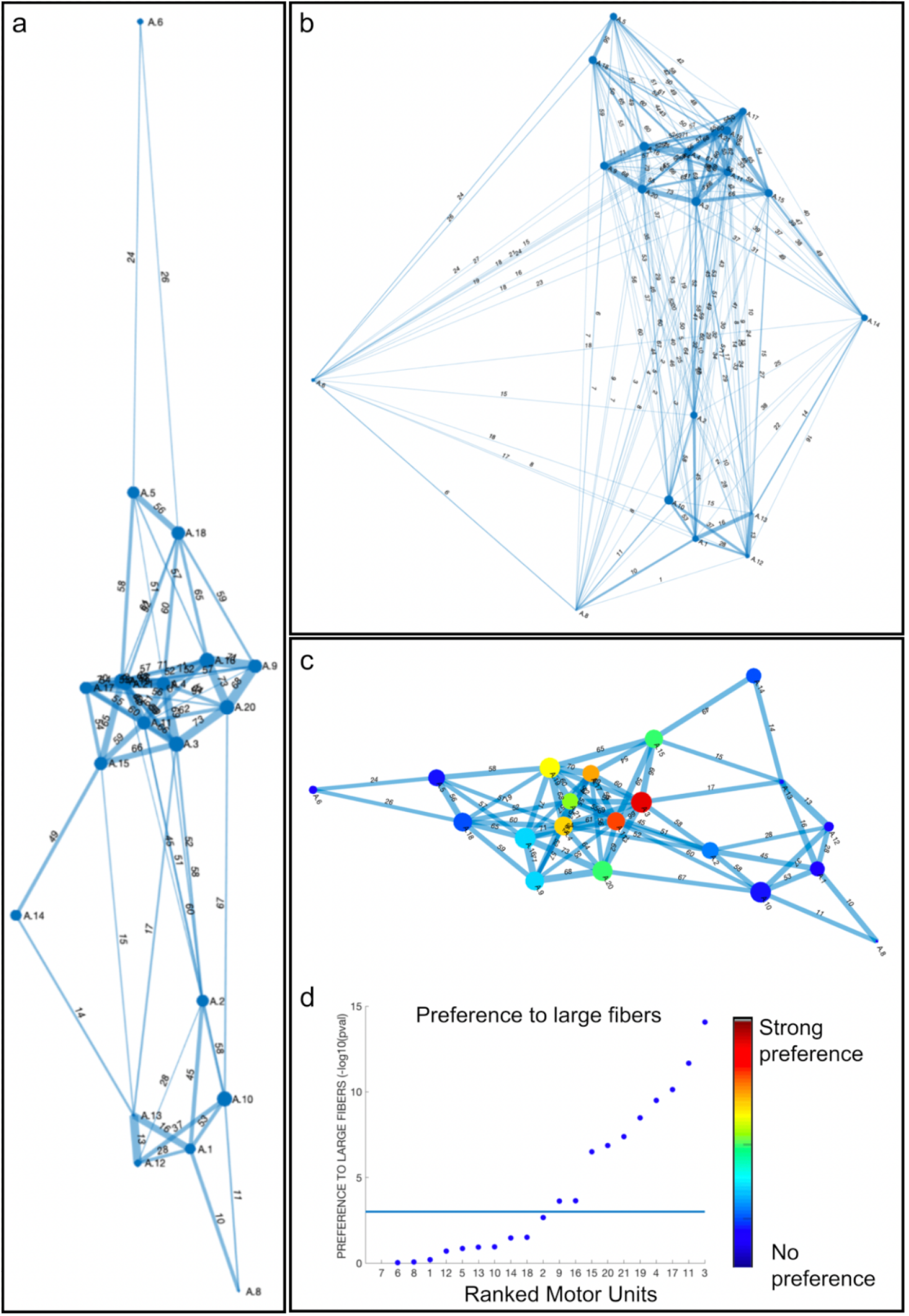
Representations of linear orders in the newborn using alternative scales. **a**, The linear orders in the paper were computed using the tendencies of pairs of axons to share more or less muscle fibers than expected by chance and without omitting edges for statistical insignificance. We repeated this analysis including only tendencies of pairs of axons to share more muscle fibers than expected by chance while only statistically significant edges are included (p<alpha=0.01). This diagram resulted in a stronger elongation than the one used for statistical analysis. **b**, The linear order was computed including all edges. Also this analysis resulted in elongated embeddings compared to embeddings of random connectomes (p<0.05). **c**, Weighting the edges as a sublinear function of -log(probability) also produces linear connectome but with less elongation compared to linear weighting (a,b). **c-d**, Ranking units based on their preference to innervate large fibers reveals that the linear order is organized according to peripheries that show no preference to large muscle fibers and a central cluster with motor units that are highly selective to large muscle fibers. This suggests that the newborn linear order is consistent with the structural properties of the muscle fibers. Moreover, even when a large motor unit, such as motor unit 10, is not selective to muscle fiber size, it is still part of the periphery of the linear order.

**Figure S5.**
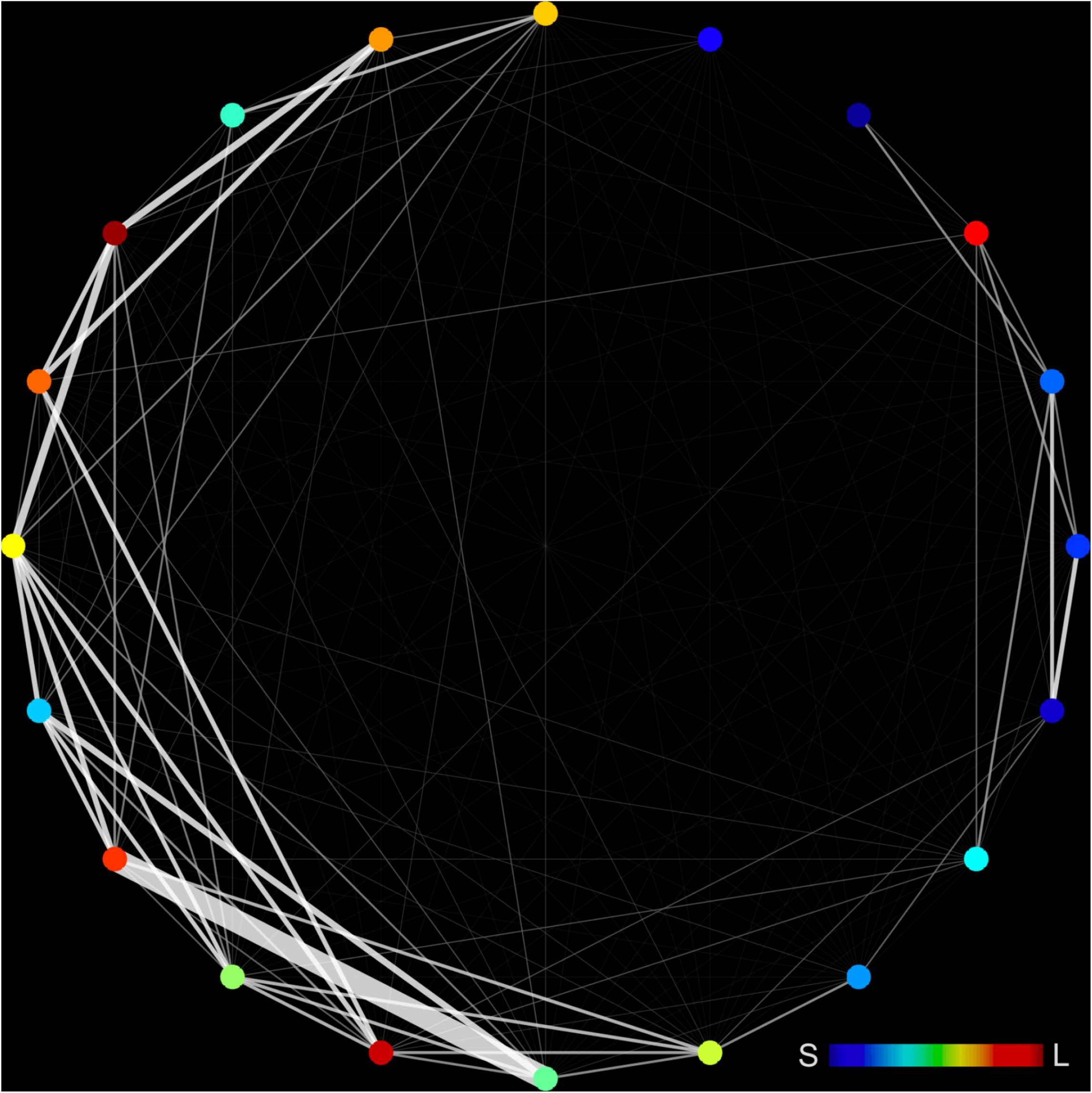
Linear order in the newborn dataset projected on a circle. A unique ranking was computed from the linear order of the newborn axons (Fig. 3a) using the coordinates of the axons along the large axis of the covariance matrix of the node’s positions. The ranking of the axons in this axis determined the location on the circle. Both edge brightness and thickness signify the probability to observe as many or more shared muscle fibers between pairs of axons as in the data compared to a random and uniform innervation model. The tendency of the thick edges to be located near the circle’s perimeter reflects the linearity of the co-innervation map.

**Figure S6.**
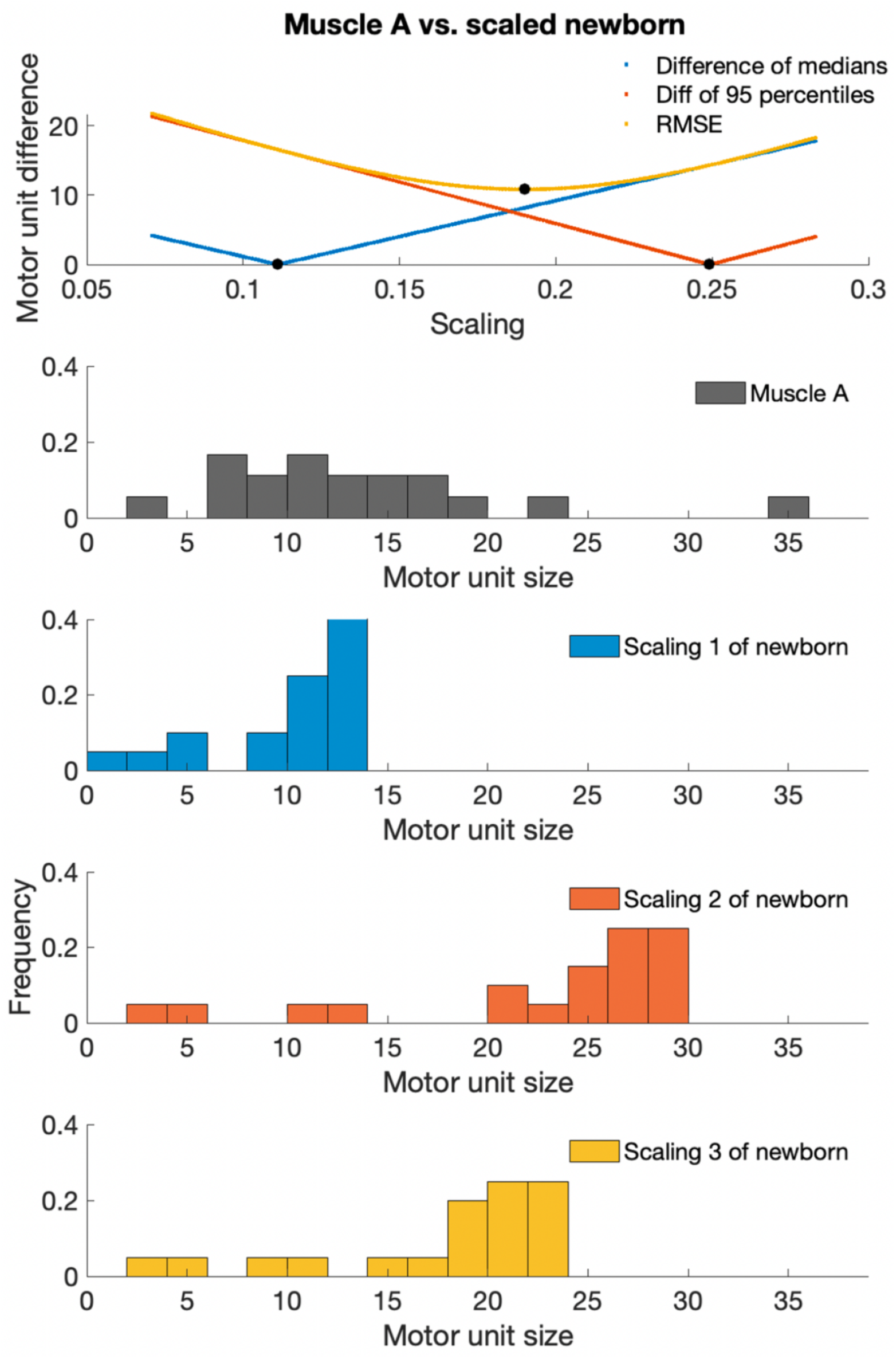
Uniform scaling of the newborn connectome. The histogram of the motor unit sizes of the newborn connectome was uniformly scaled to match that of the adult connectome (muscle A). Top: For each uniform scaling the median and 95 percentile motor unit size of the scaled-newborn and adult connectomes were compared. All possible scaling yields a large difference in the combined difference (RMSE) of the two scaling parameters. The individual histograms are presented below. The histograms of the three uniform scaling qualitatively differ from the adult histogram.

**Figure S7.**
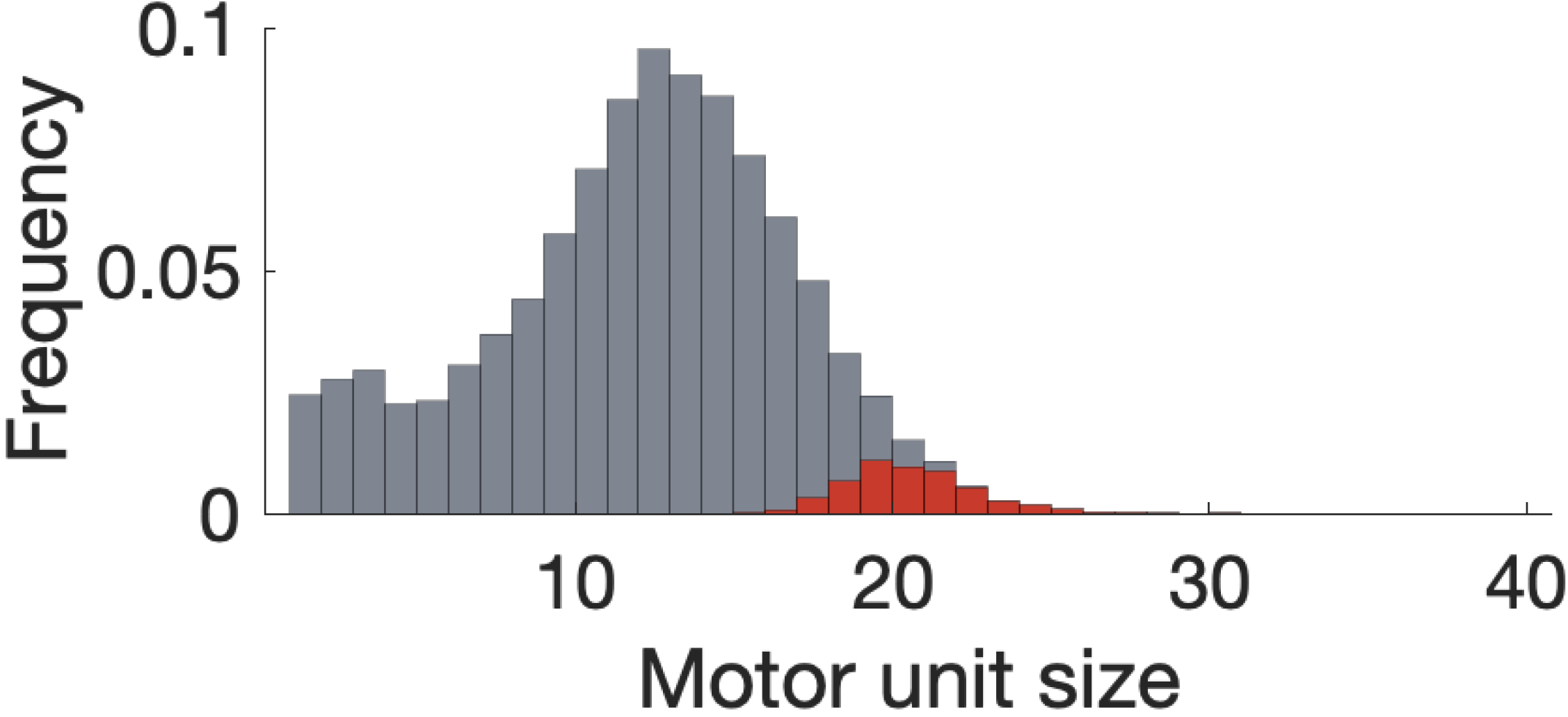
Random and gradual synapse elimination of the newborn connectome. Histogram of the sizes of motor unit sizes (gray) obtained from 500 simulated connectomes that model the process of synapse elimination. All simulated connectomes begin from the newborn connectome (Fig. 1) and connections between axons and muscle fibers are eliminated iteratively. The largest motor unit for each simulated connectome is highlighted in red. The largest motor units in the actual adult connectomes are significantly larger than the ones (n=2 and see 6 additional connectomes in (Lu et al., 2009).

**Figure S8.**
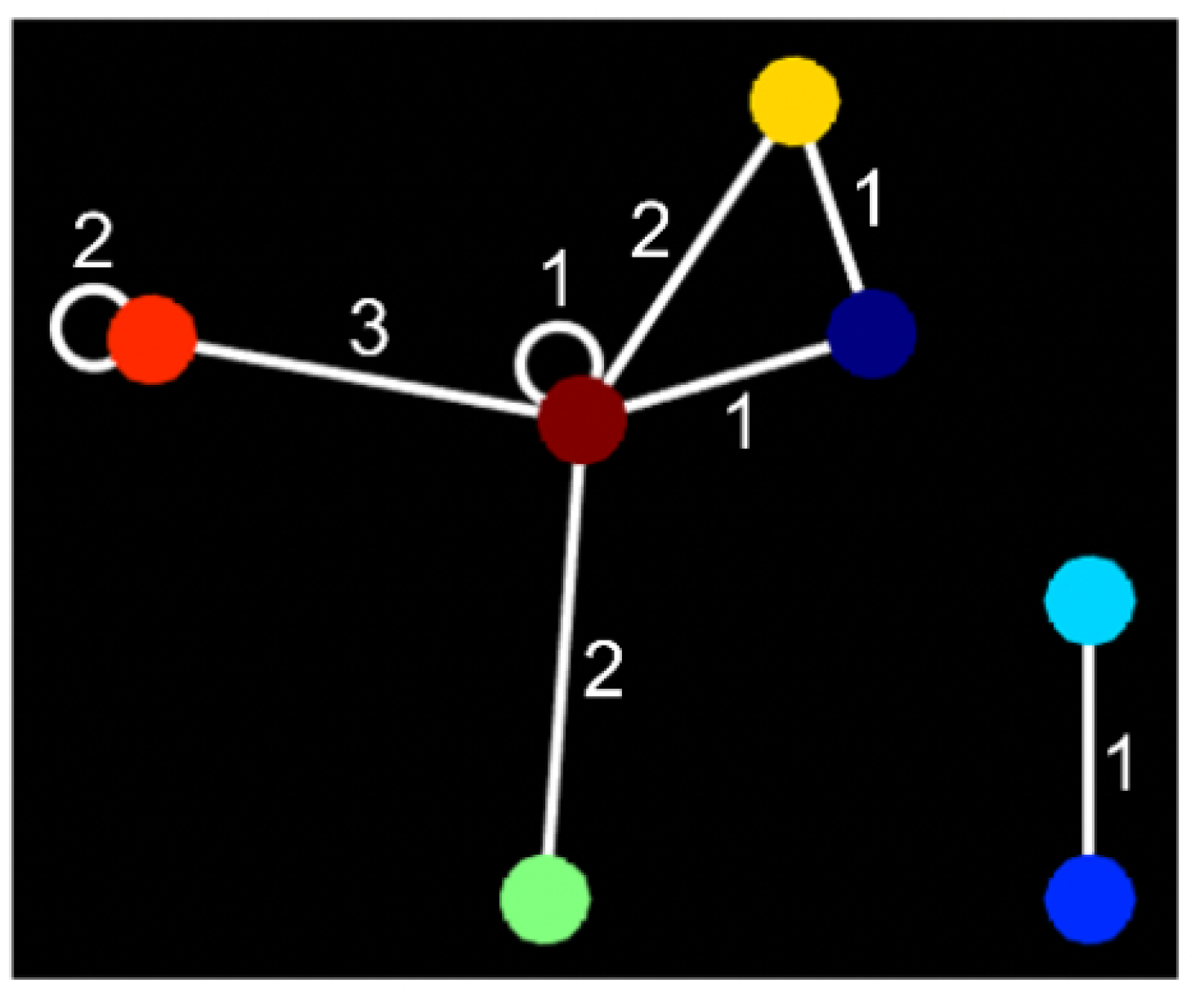
Clustered innervation on multiple-endplate fibers in the connectome of adult muscle. Seven motor units innervated 26 endplates on 13 MEFs; too few motor units to have occurred by chance (p<0.0045). The weights on the edges represent the number of distinct MEFs shared by the pair of motor units. A pair of motor units co-innervated three MEFs and two other pairs of motor units each innervated a pair of multiple-endplate muscle fibers. One motor unit (leftmost) innervated two MEFs (4 endplates). The sum of all weights corresponds to the number of endplates on MEFs.

**Figure S9.**
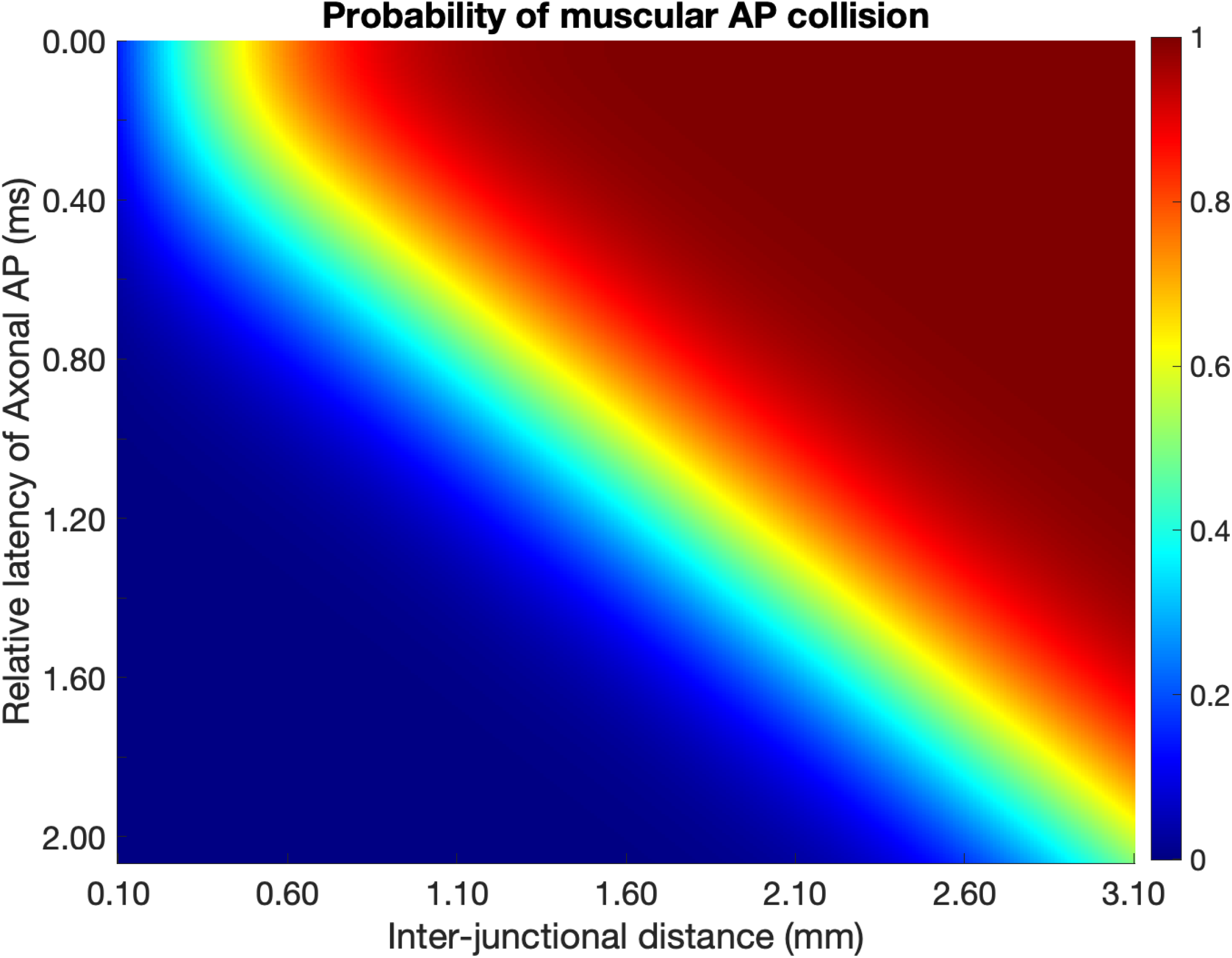
Probability of AP collision in muscle fibers depends on the inter-junctional distance and the relative latency of axonal APs. We assume that AP conduction velocity in the muscle fiber is 1.5 m/s. When the inter-junctional distance is short (< 100 μm), the probability of AP collision is low (∼ 15%) even if axonal APs arrive at the terminals without any relative latency. Therefore, one endplate can eliminate the other despite the short distance between them. On the other hand, the probability of muscular AP collision approaches 1 when the inter-junctional distance is large and the relative latency of axonal APs is not too big.

**Figure S10.**
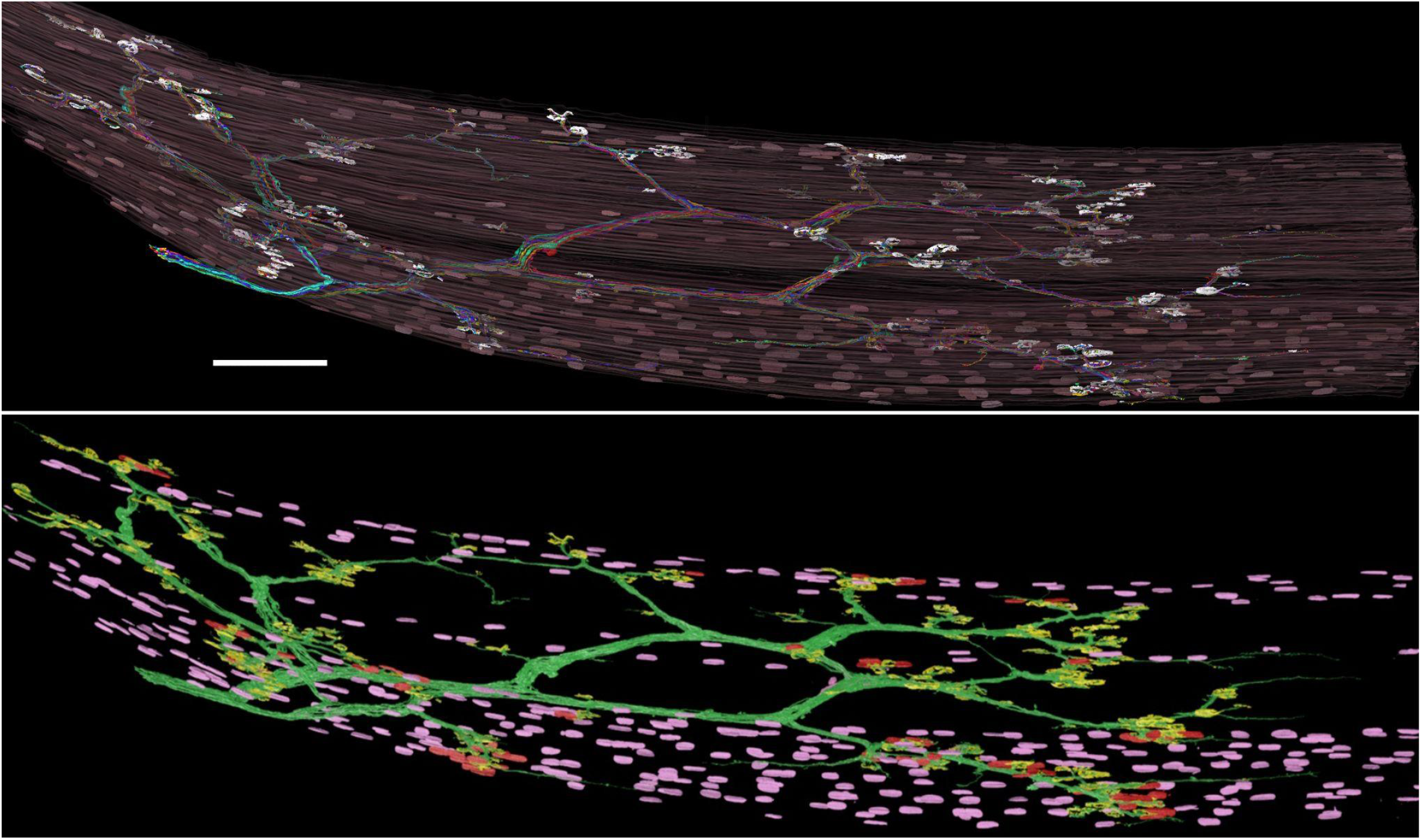
Saturation of all myonuclei in a subset of fibers. Top. All the synaptic and nonsynaptic myonuclei of 52 randomly selected muscle fibers in the newborn sample (25 MEFs and 27 SEFs), rendered with the axonal arbor, endplates and muscle fibers (top). **Bottom**. For each muscle fiber the synaptic nuclei (red) were identified in the vicinity of the endplates (yellow). The myonuclear domain of the MEFs and SEFs, which is indicative of oxidative capacity of fibers, and the number of synaptic nuclei, (Qaisar & Larsson, 2014; Van der Meer et al., 2011), were not significantly different between the two muscle fiber groups.

**Figure S11.**
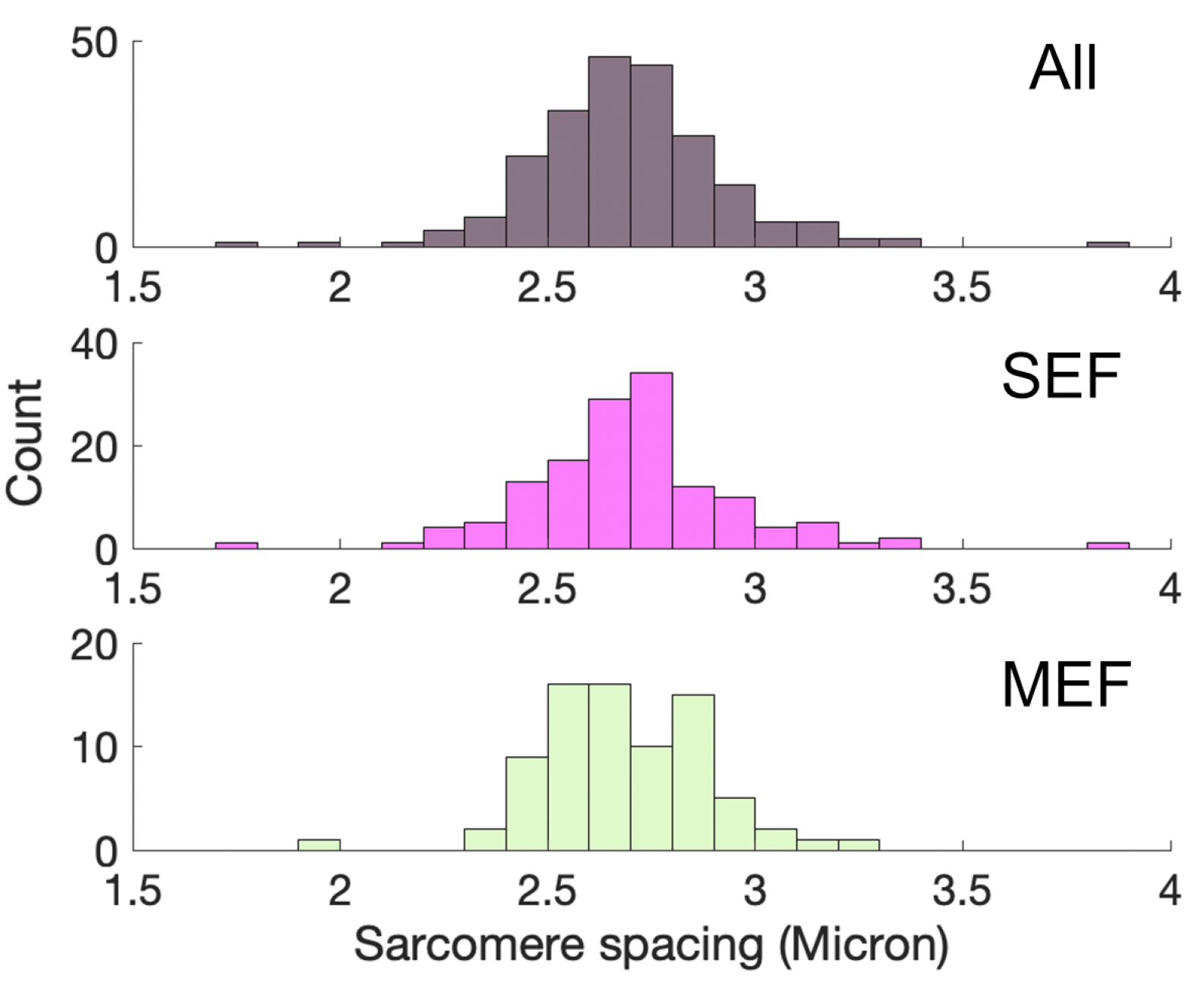
Sarcomere spacing in the newborn sample. The distribution of sarcomere spacing is unimodal and not distinctly distributed for the multiple-endplate (MEFs) and single-endplate (SEFs) fibers.

**Figure S12.**
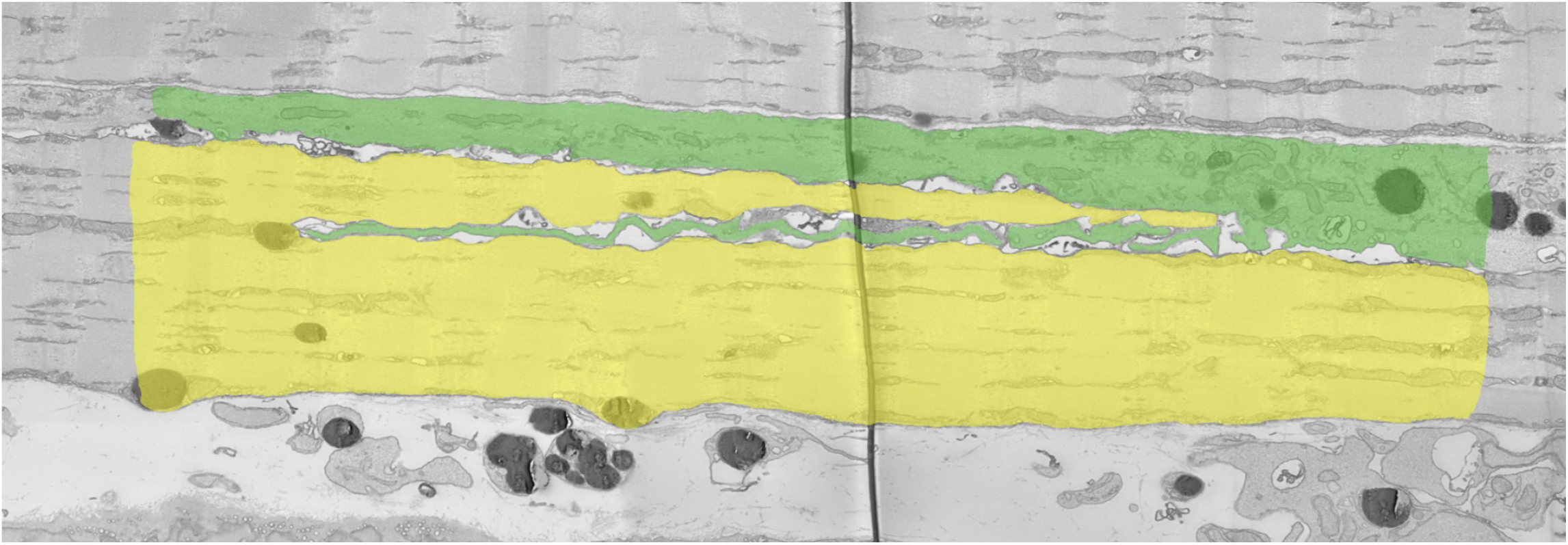
Primary and secondary fibers. In the newborn dataset we found several secondary fibers with immature contractile properties that were attached to other more mature primary fibers. In all cases the secondary fiber possessed endplates near the location of endplates on the primary fibers, as reported by (Duxson et al., 1989).

**Figure S13.**
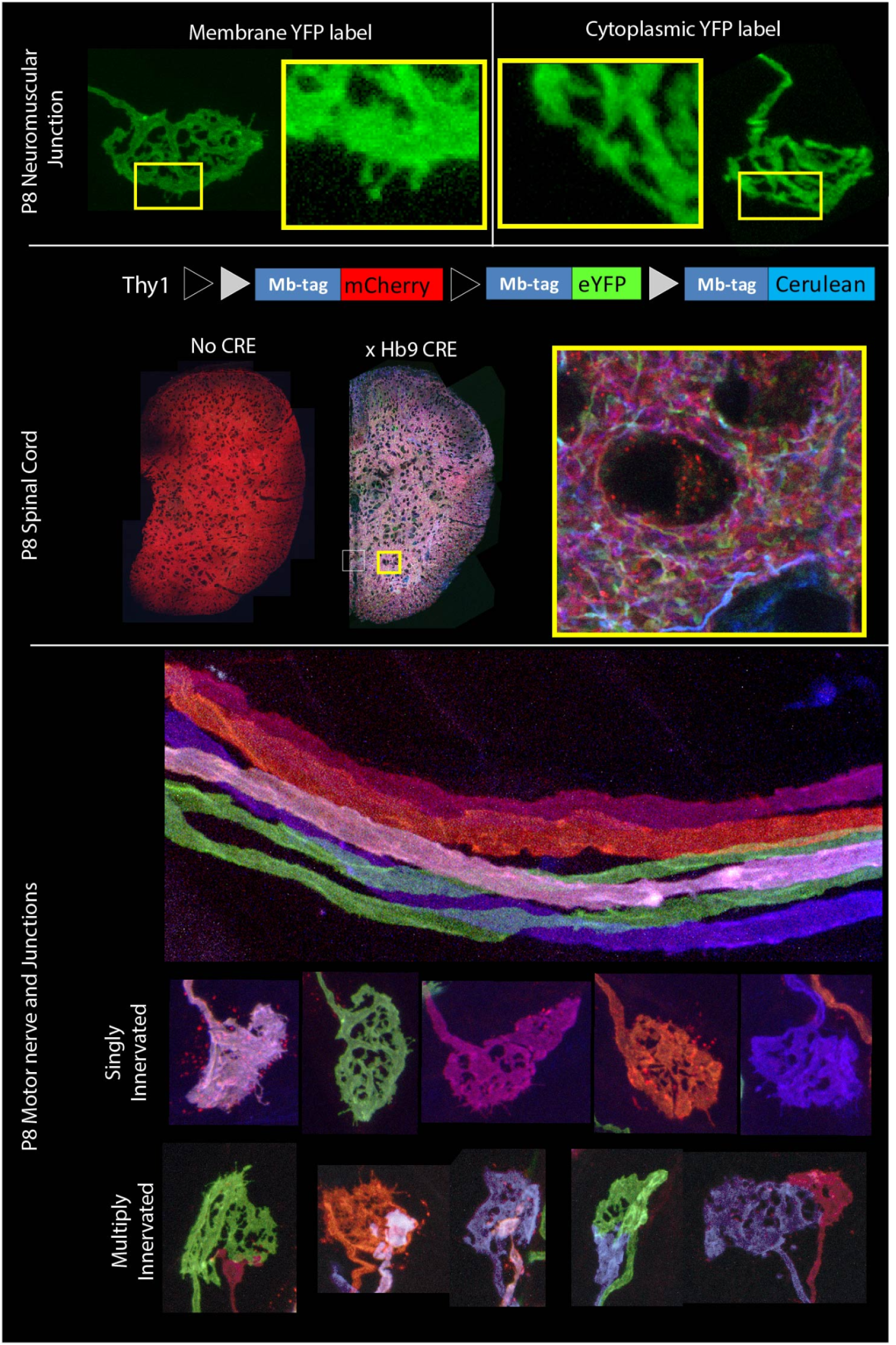
Membrane-Brainbow expression in developing motor neurons. The top panel shows a schematic of the Brainbow 1.0 type construct. Black and grey triangles indicate Lox P and 2272 sites, respectively. Blue boxes indicate the palmitolyation membrane tag (Mb-tag). The middle panel shows expression of the construct in the spinal cord with and without Cre co-expression. Yellow box shows a high resolution image of spinal cord neuropil. Bottom panel shows all the motor axons innervating a P9 Omohyiod muscle. Five unique colors and six axon collaterals are easily identifiable. Below are neuromuscular junctions from the same muscle, both singly and multiply innervated.

**Figure S14.**
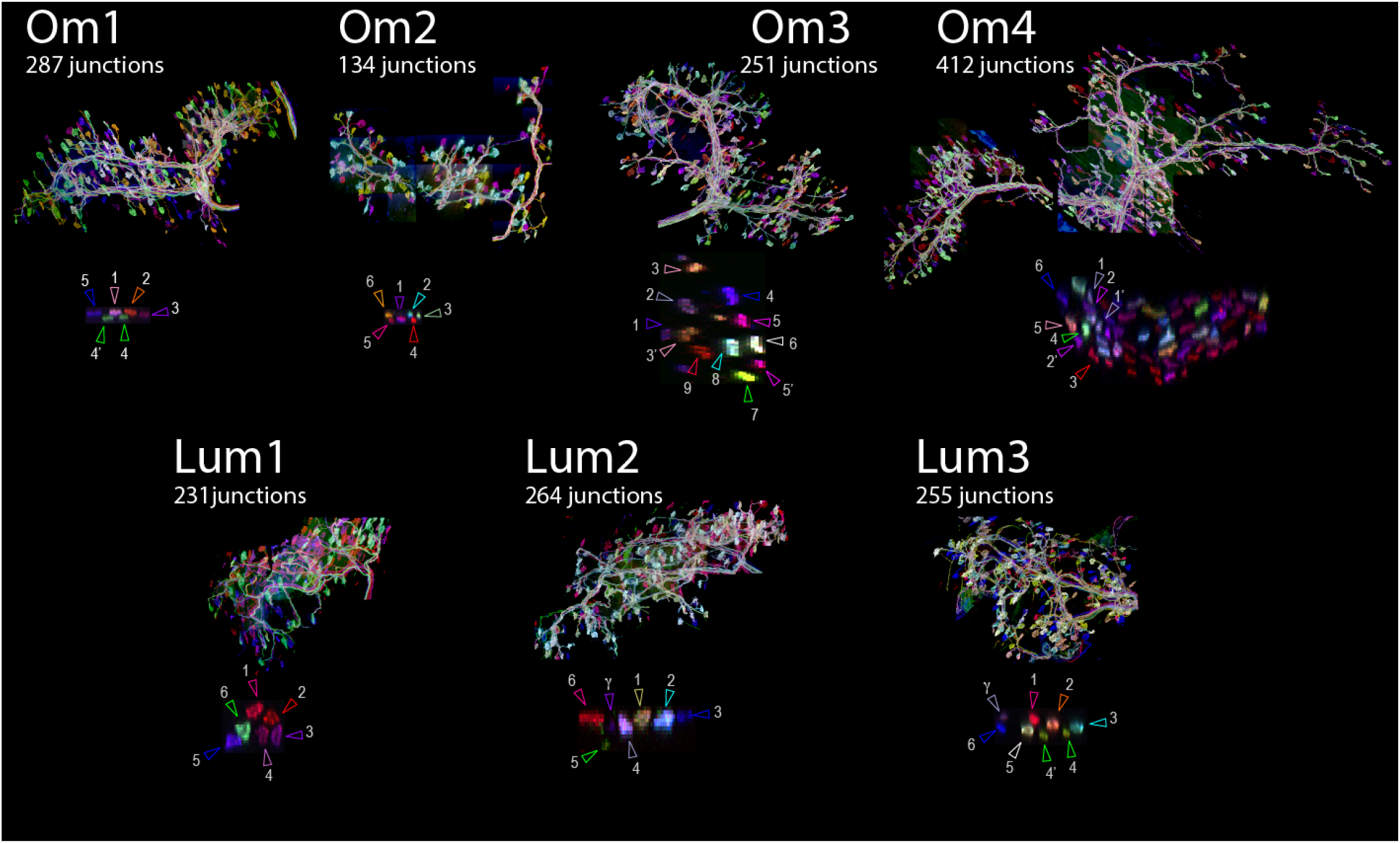
High resolution confocal image montages of brainbow labeled P6-9 Omohyoid and Lumbrical muscles. Each panel shows a downsized maximum intensity projection of a high-resolution, four channel three-dimensional image montage that contains all the motor axons and neuromuscular junctions in a developing muscle. Cross sections of the nerve at the muscle entry point are shown, and arrows identify the axons innervating the muscle (γ indicates a gamma motor axon; ’ symbol indicates a putative axon collateral (see Methods)..

**Figure S15.**
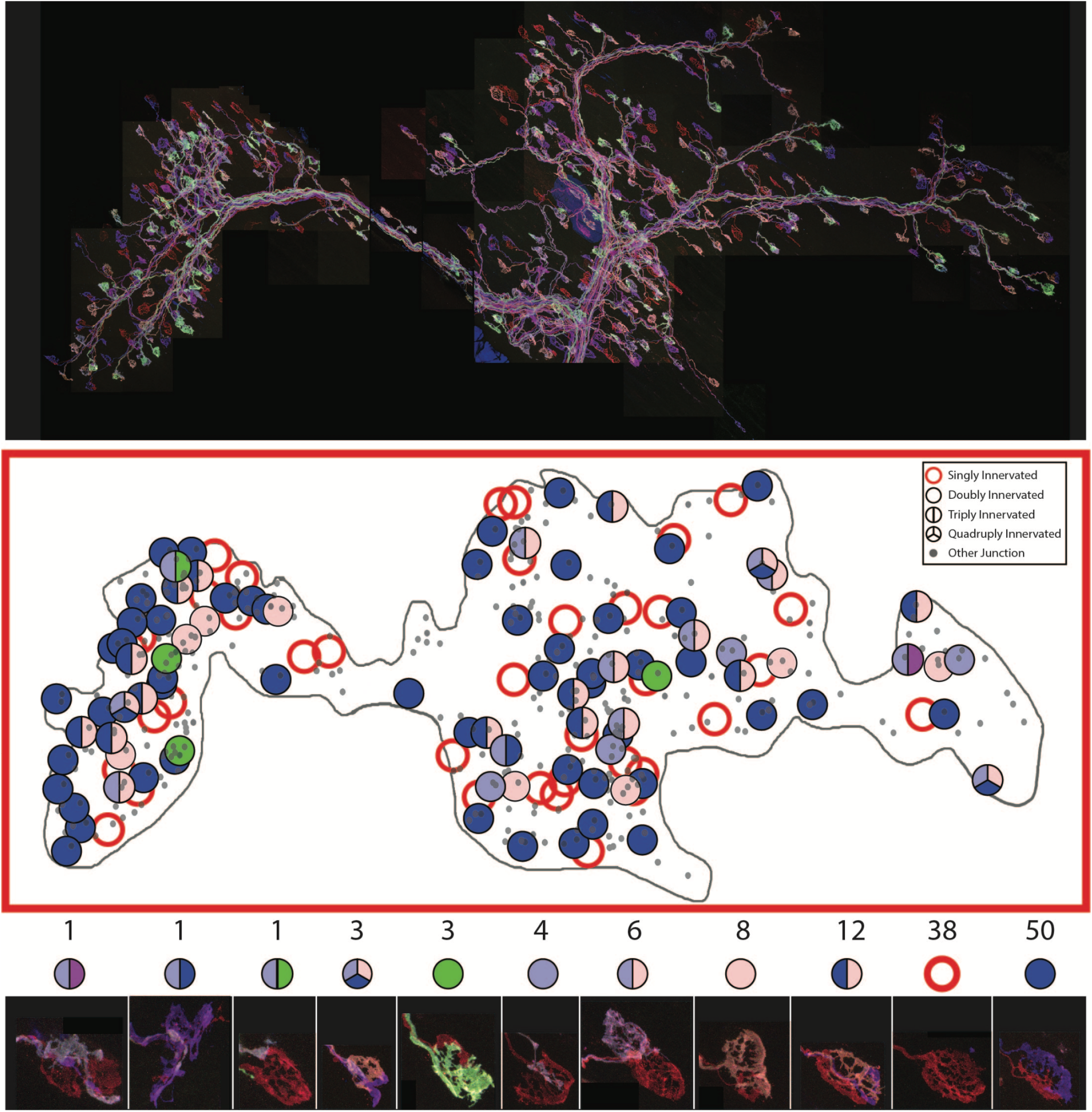
A map of all terminals of one developing motor axon. Top panel shows the maximum intensity projection of a P6 Omohyoid muscle (OM 4). The identity of all axon terminals was determined based on color. The locations of all endplates are shown as small grey dots. The locations of endplates innervated by the red-colored axon are shown as large colored circles. The color of the circles indicates the identity of other axons innervating the same endplate. The bottom panel shows images of each category of endplate as well as the numbers of endplates in each category.

**Figure S16.**
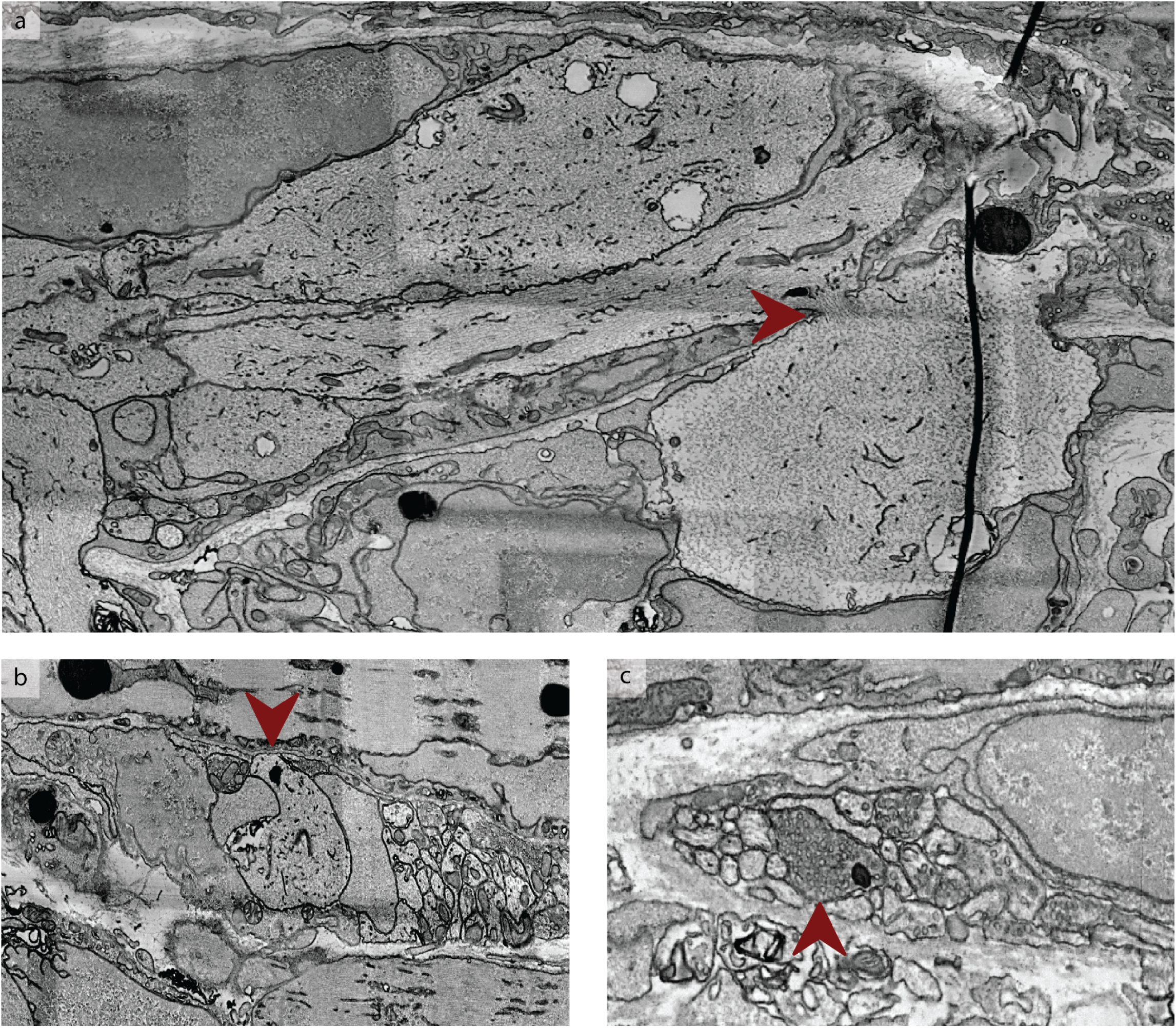
Anatomical correlates of branch-specific retraction. **a**, EM example of one axonal bulb found off of the axonal pathway. The ovoidal inflation is connected to the axon via a narrow neck (red arrowhead) where the axonal microtubules are visible. **b**, Example of putative degenerating synapse marked by the red arrow. The inflation presents signs of membrane recycling and is devoid of synaptic vesicles (present in neighboring terminals). **c**, Example of an axon bulb filled with vesicles, surrounded by axons devoid of synaptic vesicles.

**Figure S17.**
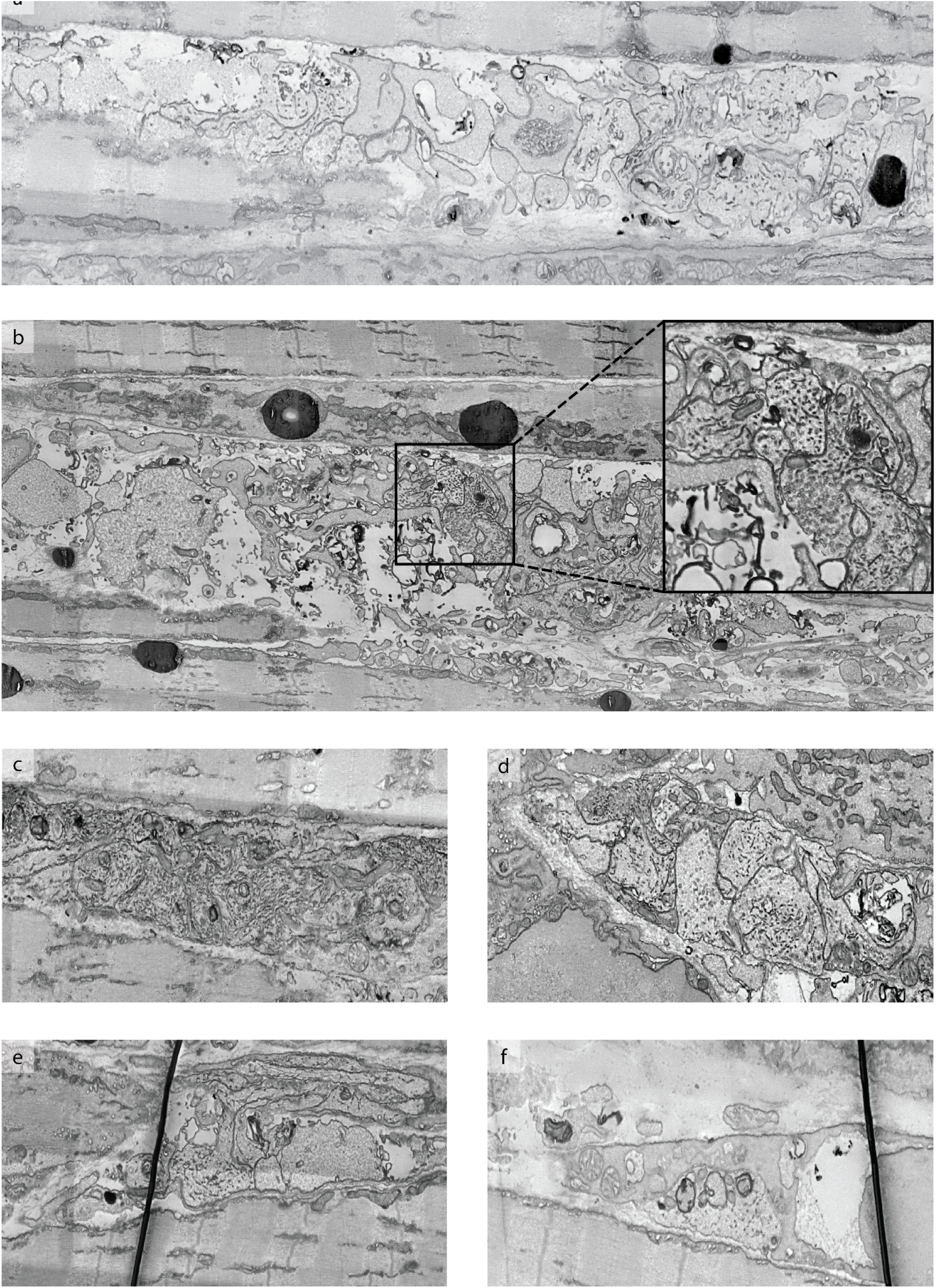
Anatomical correlates of entire sub-bundle retraction. **a, b**, Examples of graveyards. **a**, The graveyard is adjacent to non-active muscle fibers. **b**, The graveyard is near a fiber with high content of mitochondria. **c**,**d**, Examples of sub-bundles that dead-end in graveyards. Of interest are the highly concentrated black dashes in the axoplasm. **e, f**, Examples of putative degenerating synapses. Note that these structures are devoid of synaptic vesicles.

**Figure S18.**
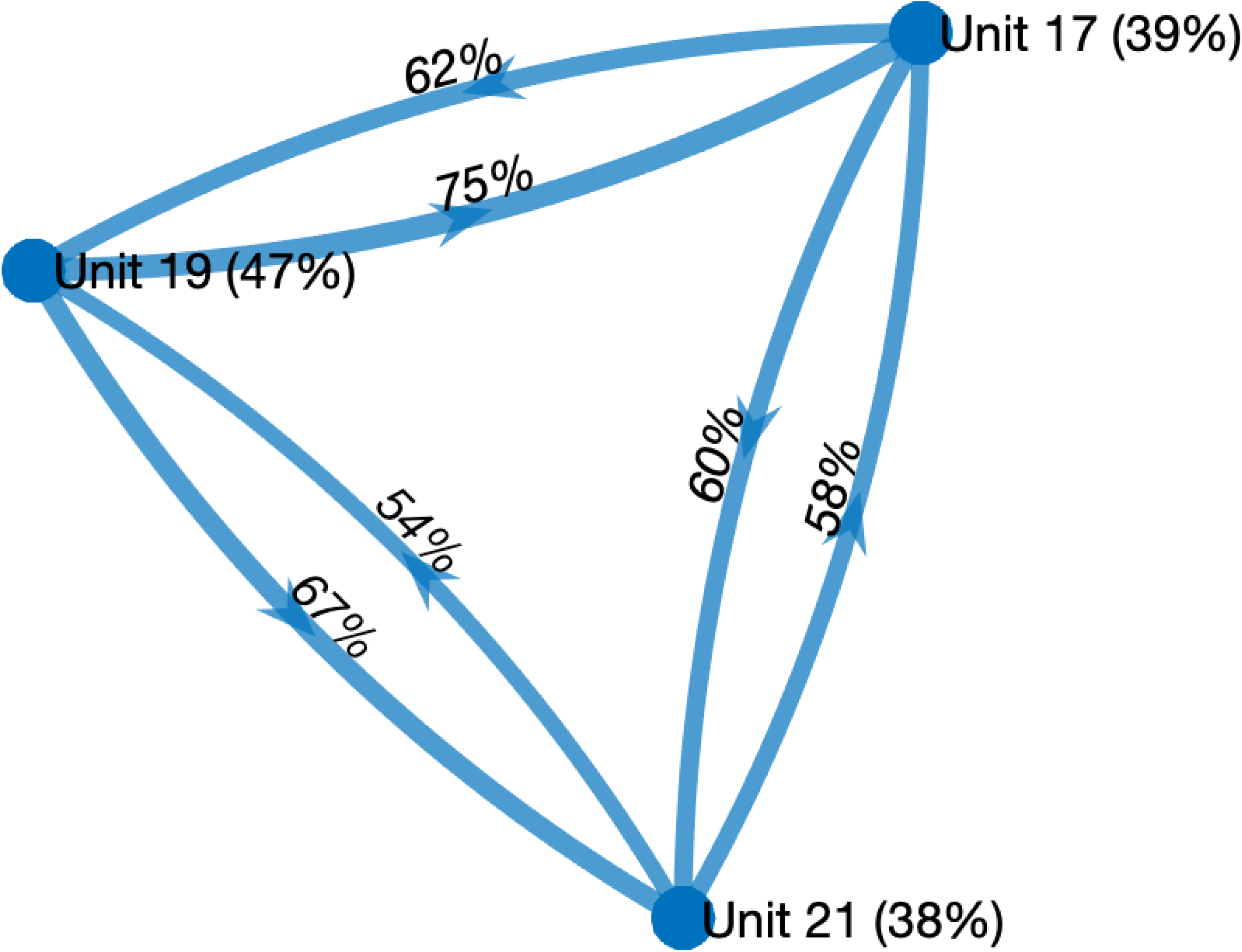
Co-innervation diagram of three motor units in the newborn dataset. Each node represents one motor unit and directed edges between two motor units (source unit and target unit) represent the proportion of muscle fibers in the target motor unit that are also included in the source motor unit. For example, unit 19 included 75% of the muscle fibers of motor unit 17. For each motor unit its size relative to the total number of fibers in the muscle (n=240) is given in parenthesis. For example, unit 17 contained 39% of the fibers in the dataset. Assuming 240 possible targets we calculate the probability to observe the number of shared targets under the assumption that targets are picked independently and uniformly at random.

## Notes

### Competing Interest Statement

The authors have declared no competing interest.

### Summary of Updates

All figures revised; abstract and introduction rewritten; author list updated.

## REFERENCES

Agduhr, E. (1919). Are the cross-striated muscle fibers of the extremities also innervated sympathetically. Proc Kon Akad Wet Amsterdam, 21, 1231–1237.

Balice-Gordon, R. J., & Lichtman, J. W. (1994). Long-term synapse loss induced by focal blockade of postsynaptlc receptors. Nature, 372(6506), 519–524.

Bendiksen, F. S., Dahl, H. A., & Teig, E. (1981). Innervation pattern of different types of muscle fibres in the human thyroarytenoid muscle. Acta Oto-Laryngologica, 91(5-6), 391–397.

Bennett, M. R. (1983). Development of neuromuscular synapses. Physiological Reviews, 63(3), 915–1048.

Bennett, M. R., & Pettigrew, A. G. (1974). The formation of synapses in striated muscle during development. In The Journal of Physiology (Vol. 241, Issue 2, pp. 515–545). https://doi.org/10.1113/jphysiol.1974.sp010670

Berger, D. R., Seung, H. S., & Lichtman, J. W. (2018). VAST (Volume Annotation and Segmentation Tool): Efficient Manual and Semi-Automatic Labeling of Large 3D Image Stacks. Frontiers in Neural Circuits, 12, 88.

Bishop, D. L., Misgeld, T., Walsh, M. K., Gan, W.-B., & Lichtman, J. W. (2004). Axon branch removal at developing synapses by axosome shedding. Neuron, 44(4), 651–661.

Brill, M. S., Kleele, T., Ruschkies, L., Wang, M., Marahori, N. A., Reuter, M. S., Hausrat, T. J., Weigand, E., Fisher, M., Ahles, A., Engelhardt, S., Bishop, D. L., Kneussel, M., & Misgeld, T. (2016). Branch-Specific Microtubule Destabilization Mediates Axon Branch Loss during Neuromuscular Synapse Elimination. Neuron, 92(4), 845–856.

Britton, T., Deijfen, M., & Martin-Löf, A. (2006). Generating simple random graphs with prescribed degree distribution. Journal of Statistical Physics, 124(6), 1377–1397.

Brown, M. C., Jansen, J. K., & Van Essen, D. (1976). Polyneuronal innervation of skeletal muscle in new-born rats and its elimination during maturation. The Journal of Physiology, 261(2), 387–422.

Buffelli, M., Busetto, G., Cangiano, L., & Cangiano, A. (2002). Perinatal switch from synchronous to asynchronous activity of motoneurons: link with synapse elimination. Proceedings of the National Academy of Sciences of the United States of America, 99(20), 13200–13205.

Burke, R. E., Levine, D. N., & Zajac, F. E., 3rd. (1971). Mammalian motor units: physiological-histochemical correlation in three types in cat gastrocnemius. Science, 174(4010), 709–712.

Busetto, G., Buffelli, M., Tognana, E., Bellico, F., & Cangiano, A. (2000). Hebbian mechanisms revealed by electrical stimulation at developing rat neuromuscular junctions. The Journal of Neuroscience: The Official Journal of the Society for Neuroscience, 20(2), 685–695.

Callaway, E. M., Soha, J. M., & Van Essen, D. C. (1987). Competition favouring inactive over active motor neurons during synapse elimination. Nature, 328(6129), 422–426.

Cattell, M. (1928). Plurisegmental innervation in the frog. The Journal of Physiology, 66(4), 431–442.

Duxson, M. J., & Sheard, P. W. (1995). Formation of new myotubes occurs exclusively at the multiple innervation zones of an embryonic large muscle. Developmental Dynamics: An Official Publication of the American Association of Anatomists, 204(4), 391–405.

Duxson, M. J., Usson, Y., & Harris, A. J. (1989). The origin of secondary myotubes in mammalian skeletal muscles: ultrastructural studies. Development, 107(4), 743–750.

Eberle, A. L., Mikula, S., Schalek, R., Lichtman, J., Tate, M. L. K., & Zeidler, D. (2015). High-resolution, high-throughput imaging with a multibeam scanning electron microscope. Journal of Microscopy, 259(2), 114–120.

Favero, M., Busetto, G., & Cangiano, A. (2012). Spike timing plays a key role in synapse elimination at the neuromuscular junction. Proceedings of the National Academy of Sciences of the United States of America, 109(25), E1667–E1675.

Favero, M., Cangiano, A., & Busetto, G. (2014). Hebb-based rules of neural plasticity: are they ubiquitously important for the refinement of synaptic connections in development? The Neuroscientist: A Review Journal Bringing Neurobiology, Neurology and Psychiatry, 20(1), 8–14.

Feng, G., Mellor, R. H., Bernstein, M., Keller-Peck, C., Nguyen, Q. T., Wallace, M., Nerbonne, J. M., Lichtman, J. W., & Sanes, J. R. (2000). Imaging neuronal subsets in transgenic mice expressing multiple spectral variants of GFP. Neuron, 28(1), 41–51.

Fischler, M. A., & Bolles, R. C. (1981). Random sample consensus: a paradigm for model fitting with applications to image analysis and automated cartography. Communications of the ACM, 24(6), 381–395.

Freund, H. J. (1983). Motor unit and muscle activity in voluntary motor control. Physiological Reviews, 63(2), 387–436.

Frontera, W. R., & Ochala, J. (2015). Skeletal muscle: a brief review of structure and function. Calcified Tissue International, 96(3), 183–195.

Fruchterman, T. M. J., & Reingold, E. M. (1991). Graph drawing by force-directed placement. Software: Practice & Experience, 21(11), 1129–1164.

Grady, R. M., Starr, D. A., Ackerman, G. L., Sanes, J. R., & Han, M. (2005). Syne proteins anchor muscle nuclei at the neuromuscular junction. Proceedings of the National Academy of Sciences of the United States of America, 102(12), 4359–4364.

Granzier, H. L., Akster, H. A., & Ter Keurs, H. E. (1991). Effect of thin filament length on the force-sarcomere length relation of skeletal muscle. The American Journal of Physiology, 260(5 Pt 1), C1060–C1070.

Hall, Z. W., & Sanes, J. R. (1993). Synaptic structure and development: the neuromuscular junction. Cell, 72 Suppl, 99–121.

Hämäläinen, N., & Pette, D. (1993). The histochemical profiles of fast fiber types IIB, IID, and IIA in skeletal muscles of mouse, rat, and rabbit. The Journal of Histochemistry and Cytochemistry: Official Journal of the Histochemistry Society, 41(5), 733–743.

Happak, W., Liu, J., Burggasser, G., Flowers, A., Gruber, H., & Freilinger, G. (1997). Human facial muscles: dimensions, motor endplate distribution, and presence of muscle fibers with multiple motor endplates. The Anatomical Record, 249(2), 276–284.

Harris, A. J., & McCaig, C. D. (1983). Embryonic proliferation of autonomic axons following muscle denervation. Neuroscience Letters, 11, S47.

Harris, W. A. (1981). Neural activity and development. Annual Review of Physiology, 43, 689–710.

Hayworth, K. J., Morgan, J. L., Schalek, R., Berger, D. R., Hildebrand, D. G. C., & Lichtman, J. W. (2014). Imaging ATUM ultrathin section libraries with WaferMapper: a multi-scale approach to EM reconstruction of neural circuits. Frontiers in Neural Circuits, 8, 68.

Henneman, E. (1957). Relation between size of neurons and their susceptibility to discharge. Science, 126(3287), 1345–1347.

Henneman, E., & Olson, C. B. (1965). RELATIONS BETWEEN STRUCTURE AND FUNCTION IN THE DESIGN OF SKELETAL MUSCLES. Journal of Neurophysiology, 28, 581–598.

Hunt, C. C., & Kuffler, S. W. (1954). Motor innervation of skeletal muscle: multiple innervation of individual muscle fibres and motor unit function. The Journal of Physiology, 126(2), 293–303.

Iwasaki, S. (1957). Mononeuronal double innervation of an amphibian striated muscle fibre. The Japanese Journal of Physiology, 7(4), 267–275.

Jarcho, L. W., Eyzaguirre, C., Berman, B., & Lilienthal, J. L., Jr. (1952). Spread of excitation in skeletal muscle; some factors contributing to the form of the electromyogram. The American Journal of Physiology, 168(2), 446–457.

Jones, S. P., Ridge, R. M., & Rowlerson, A. (1987). The non-selective innervation of muscle fibres and mixed composition of motor units in a muscle of neonatal rat. The Journal of Physiology, 386, 377–394.

Kasthuri, N., Hayworth, K. J., Berger, D. R., Schalek, R. L., Conchello, J. A., Knowles-Barley, S., Lee, D., Vázquez-Reina, A., Kaynig, V., Jones, T. R., Roberts, M., Morgan, J. L., Tapia, J. C., Seung, H. S., Roncal, W. G., Vogelstein, J. T., Burns, R., Sussman, D. L., Priebe, C. E., … Lichtman, J. W. (2015). Saturated Reconstruction of a Volume of Neocortex. Cell, 162(3), 648–661.

Katz, B., & Kuffler, S. W. (1941). Multiple motor innervation of the frog‘s Sartorius muscle. Journal of Neurophysiology, 4(3), 209–223.

Katz, B., & Miledi, R. (1965a). PROPAGATION OF ELECTRIC ACTIVITY IN MOTOR NERVE TERMINALS. Proceedings of the Royal Society of London. Series B, Containing Papers of a Biological Character. Royal Society, 161, 453–482.

Katz, B., & Miledi, R. (1965b). THE MEASUREMENT OF SYNAPTIC DELAY, AND THE TIME COURSE OF ACETYLCHOLINE RELEASE AT THE NEUROMUSCULAR JUNCTION. Proceedings of the Royal Society of London. Series B, Containing Papers of a Biological Character. Royal Society, 161, 483–495.

Krnjević, K., & Miledi, R. (1959). Presynaptic failure of neuromuscular propagation in rats. In The Journal of Physiology (Vol. 149, Issue 1, pp. 1–22). https://doi.org/10.1113/jphysiol.1959.sp006321

Lateva, Z. C., McGill, K. C., & Elise Johanson, M. (2010). The innervation and organization of motor units in a series-fibered human muscle: the brachioradialis. In Journal of Applied Physiology (Vol. 108, Issue 6, pp. 1530–1541). https://doi.org/10.1152/japplphysiol.01163.2009

Lateva, Z. C., McGill, K. C., & Johanson, M. E. (2002). Electrophysiological evidence of adult human skeletal muscle fibres with multiple endplates and polyneuronal innervation. The Journal of Physiology, 544(2), 549–565.

Lichtman, J. W., Purves, D., & Yip, J. W. (1979). On the purpose of selective innervation of guinea-pig superior cervical ganglion cells. The Journal of Physiology, 292, 69–84.

Lichtman, J. W., Purves, D., & Yip, J. W. (1980). Innervation of sympathetic neurones in the guinea-pig thoracic chain. The Journal of Physiology, 298(1), 285–299.

Lichtman, J. W., Wilkinson, R. S., & Rich, M. M. (1985). Multiple innervation of tonic endplates revealed by activity-dependent uptake of fluorescent probes. Nature, 314(6009), 357–359.

Livet, J., Weissman, T. A., Kang, H., Draft, R. W., Lu, J., Bennis, R. A., Sanes, J. R., & Lichtman, J. W. (2007). Transgenic strategies for combinatorial expression of fluorescent proteins in the nervous system. Nature, 450(7166), 56–62.

Lowe, D. G. (1999). Object recognition from local scale-invariant features. Proceedings of the Seventh IEEE International Conference on Computer Vision, 2, 1150–1157 vol.2.

Lu, J., Tapia, J. C., White, O. L., & Lichtman, J. W. (2009). The Interscutularis Muscle Connectome. In PLoS Biology (Vol. 7, Issue 2, p. e1000032). https://doi.org/10.1371/journal.pbio.1000032

McComas, A. J., Kereshi, S., & Manzano, G. (1984). Multiple innervation of human muscle fibers. Journal of the Neurological Sciences, 64(1), 55–64.

Meister, M., Wong, R. O., Baylor, D. A., & Shatz, C. J. (1991). Synchronous bursts of action potentials in ganglion cells of the developing mammalian retina. Science, 252(5008), 939–943.

Morgan, J. L., Berger, D. R., Wetzel, A. W., & Lichtman, J. W. (2016). The Fuzzy Logic of Network Connectivity in Mouse Visual Thalamus. Cell, 165(1), 192–206.

Newman, M. (2018). Networks. Oxford University Press.

Njå, A., & Purves, D. (1977). Specific innervation of guinea-pig superior cervical ganglion cells by preganglionic fibres arising from different levels of the spinal cord. The Journal of Physiology, 264(2), 565–583.

Pavarino, E., Berger, D. R., Morozova, O., Drescher, B., Bidel F., Kang K., Lichtman J. W., Meirovitch Y. (2019). mEMbrain: an interactive deep learning tool for labeling and instance segmentation. Bernstein Conference for Computational Neuroscience, Berlin.

Périé, S., St Guily, J. L., Callard, P., & Sebille, A. (1997). Innervation of adult human laryngeal muscle fibers. Journal of the Neurological Sciences, 149(1), 81–86.

Personius, K. E., & Balice-Gordon, R. J. (2001). Loss of correlated motor neuron activity during synaptic competition at developing neuromuscular synapses. Neuron, 31(3), 395–408.

Preibisch, S., Saalfeld, S., & Tomancak, P. (2009). Globally optimal stitching of tiled 3D microscopic image acquisitions. Bioinformatics, 25(11), 1463–1465.

Purves, D., & Lichtman, J. W. (1980). Elimination of synapses in the developing nervous system. Science, 210(4466), 153–157.

Qaisar, R., & Larsson, L. (2014). What determines myonuclear domain size? Indian Journal of Physiology and Pharmacology, 58(1), 1–12.

Redfern, P. A. (1970). Neuromuscular transmission in new-born rats. The Journal of Physiology, 209(3), 701–709.

Riley, D. A. (1981). Ultrastructural evidence for axon retraction during the spontaneous elimination of polyneuronal innervation of the rat soleus muscle. Journal of Neurocytology, 10(3), 425–440.

Rossi, G., & Cortesina, G. (1965). MORPHOLOGICAL STUDY OF THE LARYNGEAL MUSCLES IN MAN. INSERTIONS AND COURSES OF THE MUSCLE FIBRES, MOTOR END-PLATES AND PROPRIOCEPTORS. Acta Oto-Laryngologica, 59, 575–592.

Rublee, E., Rabaud, V., Konolige, K., & Bradski, G. (2011). ORB: An efficient alternative to SIFT or SURF. 2011 International Conference on Computer Vision, 2564–2571.

Saalfeld, S., Fetter, R., Cardona, A., & Tomancak, P. (2012). Elastic volume reconstruction from series of ultra-thin microscopy sections. Nature Methods, 9(7), 717–720.

Sandmann, G. (1885). Ueber die Vertheilung der motorischen Nervenendapparate in den quergestreiften Muskeln der Wirbelthiere. Virchows Archiv Fur Pathologische Anatomie Und Physiologie Und Fur Klinische Medizin, 240–252.

Sanes, J. R., & Lichtman, J. W. (1999). Development of the vertebrate neuromuscular junction. Annual Review of Neuroscience, 22, 389–442.

Sanes, J. R., & Lichtman, J. W. (2001). Induction, assembly, maturation and maintenance of a postsynaptic apparatus. Nature Reviews. Neuroscience, 2(11), 791–805.

Schindelin, J., Arganda-Carreras, I., Frise, E., Kaynig, V., Longair, M., Pietzsch, T., Preibisch, S., Rueden, C., Saalfeld, S., Schmid, B., Tinevez, J.-Y., White, D. J., Hartenstein, V., Eliceiri, K., Tomancak, P., & Cardona, A. (2012). Fiji: an open-source platform for biological-image analysis. Nature Methods, 9(7), 676–682.

Stryker, M. P., & Harris, W. A. (1986). Binocular impulse blockade prevents the formation of ocular dominance columns in cat visual cortex. The Journal of Neuroscience: The Official Journal of the Society for Neuroscience, 6(8), 2117–2133.

Tapia, J. C., Kasthuri, N., Hayworth, K. J., Schalek, R., Lichtman, J. W., Smith, S. J., & Buchanan, J. (2012). High-contrast en bloc staining of neuronal tissue for field emission scanning electron microscopy. Nature Protocols, 7(2), 193–206.

Tapia, J. C., Wylie, J. D., Kasthuri, N., Hayworth, K. J., Schalek, R., Berger, D. R., Guatimosim, C., Seung, H. S., & Lichtman, J. W. (2012). Pervasive synaptic branch removal in the mammalian neuromuscular system at birth. Neuron, 74(5), 816–829.

Tseng, B. S., Kasper, C. E., & Edgerton, V. R. (1994). Cytoplasm-to-myonucleus ratios and succinate dehydrogenase activities in adult rat slow and fast muscle fibers. Cell and Tissue Research, 275(1), 39–49.

Tsuriel, S., Gudes, S., Draft, R. W., Binshtok, A. M., & Lichtman, J. W. (2015). Multispectral labeling technique to map many neighboring axonal projections in the same tissue. Nature Methods, 12(6), 547–552.

Turney, S. G., & Lichtman, J. W. (2012). Reversing the outcome of synapse elimination at developing neuromuscular junctions in vivo: evidence for synaptic competition and its mechanism. PLoS Biology, 10(6), e1001352.

Van der Meer, S. F. T., Jaspers, R. T., & Degens, H. (2011). Is the myonuclear domain size fixed? Journal of Musculoskeletal & Neuronal Interactions, 11(4), 286–297.

Walker, L. B., Jr. (1961). Multiple motor innervation of individual muscle fibers in the M. tibialis anterior of the dog. The Anatomical Record, 139, 1–12.

Wang, M., Zhao, Y., & Zhang, B. (2015). Efficient Test and Visualization of Multi-Set Intersections. Scientific Reports, 5, 16923.

Willshaw, D. J. (1981). The establishment and the subsequent elimination of polyneuronal innervation of developing muscle: theoretical considerations. Proceedings of the Royal Society of London. Series B, Containing Papers of a Biological Character. Royal Society, 212(1187), 233–252.

Witvliet, D., Mulcahy, B., Mitchell, J. K., Meirovitch, Y., Berger, D. R., Wu, Y., Liu, Y., Koh, W. X., Parvathala, R., Holmyard, D., Schalek, R. L., Shavit, N., Chisholm, A. D., Lichtman, J. W., Samuel, A. D. T., & Zhen, M. (2021). Connectomes across development reveal principles of brain maturation. Nature, 596(7871), 257–261.

Wyatt, R. M., & Balice-Gordon, R. J. (2003). Activity-dependent elimination of neuromuscular synapses. Journal of Neurocytology, 32(5-8), 777–794.

Zajac, F. E., & Faden, J. S. (1985). Relationship among recruitment order, axonal conduction velocity, and muscle-unit properties of type-identified motor units in cat plantaris muscle. Journal of Neurophysiology, 53(5), 1303–1322.

Zenker, W., Snobl, D., & Boetschi, R. (1990). Multifocal innervation and muscle length. Anatomy and Embryology, 182(3), 273–283.

Zucker, R. S. (1973). Theoretical implications of the size principle of motoneurone recruitment. Journal of Theoretical Biology, 38(3), 587–596.

